# Engineered HMGB1 construct with tandem Box B domains promotes tissue regeneration without potential for inflammation

**DOI:** 10.1101/2024.11.06.622383

**Authors:** Álvaro Viñals Guitart, Carl Lee, Ana Isabel Espírito Santo, Jia-Ling Ruan, Shih-Hsuan Mao, Lynn William, Christina Redfield, Oleg Fedorov, Nicola A Burgess-Brown, Wyatt W. Yue, Jagdeep Nanchahal

**Affiliations:** Centre for Medicines Discovery, University of Oxford, Oxford, UK; Kennedy Institute of Rheumatology, University of Oxford, Oxford, UK; Department of Oncology, University of Oxford, Oxford, UK; Department of Plastic and Reconstructive Surgery, Chang Gung Memorial Hospital, Linkuo & Chang Gung University College of Medicine, Taiwan; Department of Biochemistry, University of Oxford, Oxford, UK; Bioscience Institute, Newcastle University, Newcastle upon Tyne, UK; Pharmaceutical & Biological Chemistry, UCL, London, UK

## Abstract

Fully reduced High Mobility Group Box 1 (HMGB1) binds CXCL12 and signals via CXCR4 when released into the extracellular space. It acts as a chemokine and transitions stem cells from quiescent G₀ to a primed G_Alert_ state. Cells in G_Alert_ rapidly enter G_1_ in response to activating factors released by tissue injury to promote tissue repair. However, oxidative conversion of FR-HMGB1 into the disulfide form activates proinflammatory pathways via TLR-2, TLR-4 and RAGE. Peptide mapping and NMR spectroscopy identified a conserved CXCL12-binding motif within each Box and adjacent flanking regions. We decoupled the regenerative and inflammatory functions using an engineered construct (dBB12L), comprising tandem Box B domains with a flexible linker. dBB12L exhibited CXCL12 binding and accelerated repair equivalent to FR-HMGB1. Importantly, dBB12L lacked detectable RAGE binding and did not signal via TLR-2 and TLR-4, establishing it as a potential therapeutic to promote tissue repair without deleterious inflammation.

## INTRODUCTION

Resident stem and progenitor cells play a key role in homeostasis and repair following injury^1^. However, in most adult tissues, healing occurs predominantly through scarring^2^ rather than regeneration. Whilst exogenous administration of stem cells is now standard of care for some hematological disorders^3^, similar efforts to use stem cells to regenerate solid organs have largely failed^4^. This is due, in part, to the inflammatory environment and the subsequent fibrosis following injury, which hinders stem cell engraftment a n d disrupts the stem cell niche^5^. Therefore, focus has shifted to promoting tissue regeneration by activating endogenous regenerative repair processes^6–8^. Development of a successful therapeutic would depend on identification of soluble mediators to promote these pathways^9^.

We previously identified High Mobility Group Box 1 (HMGB1) as a key mediator of regeneration across multiple tissues, including bone, blood and skeletal muscle^10^. HMBG1 is composed of two DNA-binding domains, Box A (8-78) and Box B (94-162), each comprising three lysine and arginine-rich α-helices (I-III), arranged in an L-shaped conformation. These domains are connected by a flexible linker and followed by a negatively charged acidic tail (Figure S1). *In vivo*, HMGB1 exists in three redox states: fully reduced (FR-HMGB1), disulfide (DS-HMGB1) with a disulfide bond between C22 and C44 in Box A, and a fully oxidized inactive sulfonyl form (SO3-HMGB1) where C22, C44 and C105 are terminally oxidized to sulfonate^11^.

FR-HMGB1, the most abundant non-histone nuclear protein, regulates a variety of processes, including gene transcription and DNA repair^12,13^. It also functions extracellularly as a prototypical alarmin, a constitutive cell component triggering immune activity when released from cells^14^. While extracellular HMGB1 is often considered proinflammatory^15^, this activity is only associated with the DS-HMGB1 form. In contrast, FR-HMGB1 promotes tissue regeneration. Following cell injury or non-programmed death^11^, FR-HMGB1 is released and binds to the chemokine CXCL12, forming a heterocomplex that activates the CXCR4 receptor. This interaction drives stem and progenitor cells into G_Alert_^10^,an intermediate phase between G_0_ and G_1_. When exposed to appropriate activating factors, cells in G_Alert_ rapidly enter G_1_ to effect tissue repair^16^. However, FR-HMGB1 is susceptible to oxidation in the inflammatory environment^17,18^, with a conversion half-life o f a b o u t 17 min in human fluids in vitro^19^ . This conversion to DS-HMGB1 transforms HMGB1 from a regenerative signal to a proinflammatory mediator. Additionally, DS-HMGB1 is actively secreted by immune cells in response to inflammatory stimuli^20,21^. DS-HMGB1 signals through receptors such as TLR-2^22–24^, TLR-4/MD-2 complexes^23,25^ and RAGE^26,27^, converging on MyD88^28,29^ and activating NF-κβ pathways. These pathways not only exacerbate inflammation but also impair tissue repair^30–32^. DS-HMGB1 is also an important activator of platelets via RAGE^33^, increasing the risk of thrombosis.

The therapeutic use of native FR-HMGB1 is limited by its susceptibility to oxidation in the acidic milieu of injured tissues, which leads to rapid conversion into DS-HMGB1. 3S-HMGB1, where the three critical cysteine residues (C22, C44, and C105) have been substituted with serine residues has been described^11^. This engineered HMGB1 variant is resistant to oxidation, does not bind TLR-4, and retains regenerative function in skeletal muscle, bone and haematological injury models^34,35^. However, the broader immunological signaling profile of 3S-HMGB1 remains incompletely understood, particularly with respect to its ability to engage other key inflammatory receptors such as RAGE and TLR-2. Importantly, in the context of myocardial infarction, 3S-HMGB1 promoted fibrosis, whereas native FR-HMGB1 promoted functional recovery^36^, highlighting the context-specific consequences of modulating the receptor interactions of HMGB1.

Therefore, to advance the therapeutic utility of HMGB1-based strategies, it is critical to develop variants that retain pro-reparative functions across all tissues by preserving its ability to form a heterocomplex with CXCL12 and activate CXCR4 signaling, while eliminating potential for pro-inflammatory signaling. Here, we report the development of an engineered HMGB1 construct (dBB12L) designed to preserve the full regenerative potential of FR-HMGB1 while eliminating binding to RAGE, TLR-2 and TLR-4/MD-2. We characterize the biochemical and structural properties of this construct and demonstrate its ability to activate endogenous repair pathways across multiple tissue models without triggering inflammatory responses. This positions dBB12L as a potential therapeutic to promote repair of multiple tissues.

## RESULTS

### CXCL12 binding regions are conserved across both Box domains and adjacent flanking regions of HMGB1

We mapped the regions of HMGB1 involved in CXCL12 binding with spot arrays using overlapping peptides spanning the full HMGB1 sequence probed with recombinant CXCL12-His6. This approach revealed two discrete CXCL12-binding regions within each Box domain (Fig. 1a). The first was located within α-helix I and extended into the N-terminal segment of α-helix II, overlapping the known glycyrrhizin binding site⁴¹. This first set of peptides covered residues Asp4–Ser41 within Box A and Asp90–Gly122 within Box B; in both domains this overlap coincides with the glycyrrhizin-binding region that inhibits the regenerative effects of the HMGB1–CXCL12 heterocomplex^37^. Interestingly, this binding region also extends beyond the UniProt-defined structural boundaries of Box A (Pro8–Ile78) and Box B (Pro94–Arg162), incorporating part of the N-terminus not represented in our initial Box A and Box B constructs. The second CXCL12-binding region mapped to the C-terminal end of α-helix III, spanning Lys65–Gly82 in Box A and Lys151–Asp168 in Box B, with the signal intensity in Box B less than for Box A.

**Fig. 1:**
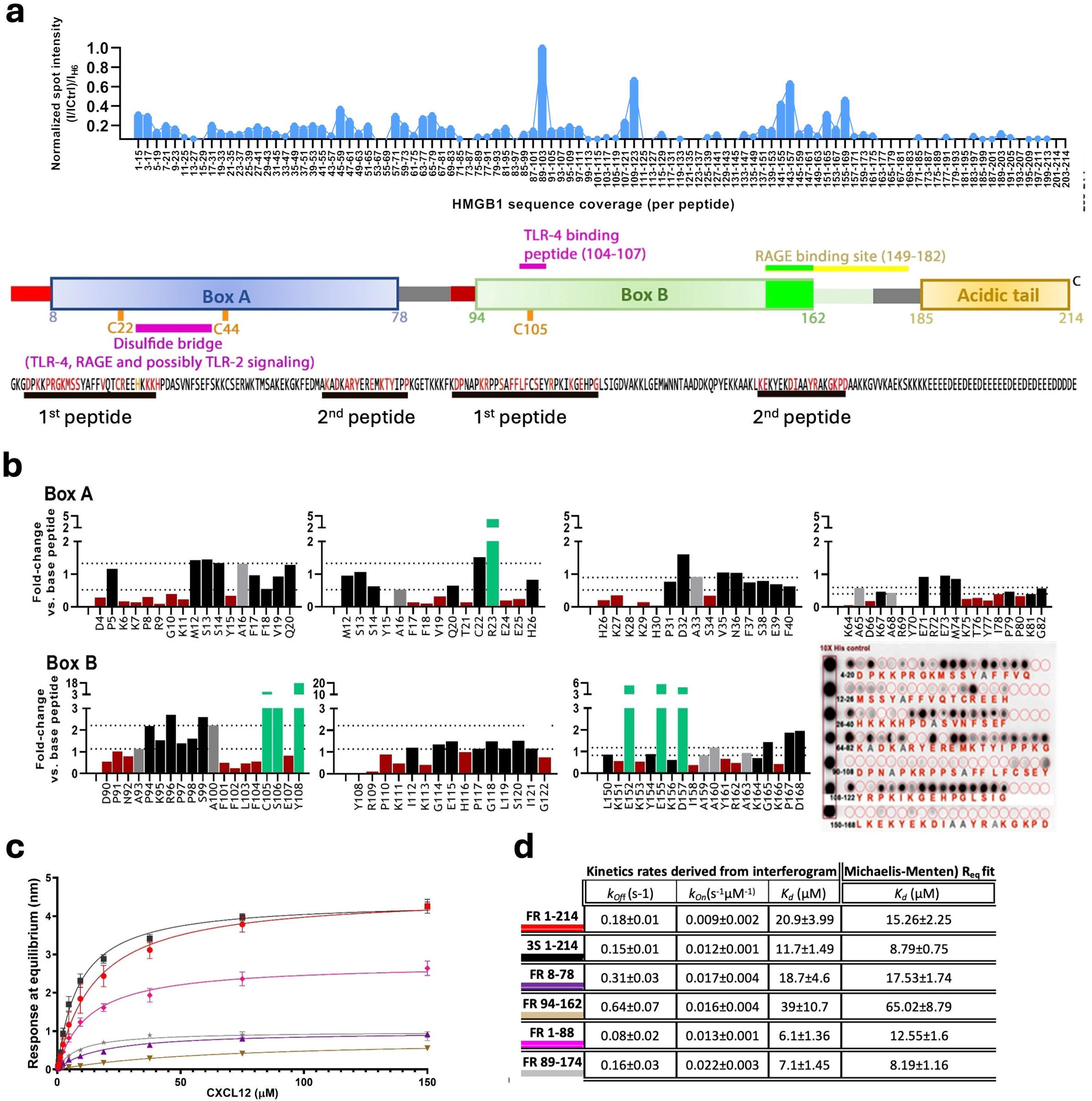
Residues critical for CXCL12 binding are conserved across both Box domains. **(a)** Top: Intensity densitogram for peptide array of HMGB1 15-mer peptides probed with CXCL12-His6. Bottom: Schematic of FL-HMGB1 with sequence. CXCL12 binding peptides are underlined in black, and key residues observed in the alanine scan are highlighted in red. **(b)** Quantification of spot intensity of alanine scan of the CXCL12 binding peptides identified in the control array normalized to the unmodified peptide control. Red indicates diminished, green increased binding to CXCL12. Alanine residues (no mutation) are shown in grey as variation threshold. A membrane developed with ECL is shown with the corresponding residues mutated to alanine. **(c)** Michaelis-Menten saturation fits from BLI interferograms of CXCL12 binding to biotinylated HMGB1 constructs. Color of individual HMGB1 construct is highlighted on the right with the summary of kinetic parameters. **(d)** *K_d_* values across constructs were compared using post-hoc 2-way ANOVA when AICc supported a model with multiple constants. R_eq_ (response at equilibrium), *k_Off_* [dissociation constant (s^-1^)], *k_On_* [association constant (µM*s)^-1^].

Next, individual Ala scanning substitutions within these peptide sequences identified key residues flanking the Box domains as critical for interaction with CXCL12 (Fig. 1b). Specifically, the first 4 residues before each Box (D4, K6, K7 N-terminal to Box A, and D90, N92 N-terminal to Box B) and the C-terminal residues (I78-P80 C-terminal to Box A and K166 in C-terminal to Box B) were found to substantially contribute to CXCL12 binding.

We validated these results by performing a reverse peptide array in which CXCL12-derived peptides were probed with full-length HMGB1 proteins. Both FR-HMGB1 and the non-oxidisable 3S-HMGB1 (1–214) bound to CXCL12 sequences spanning the entire β-sheet region (highlighted in red and blue, Figure S2). In contrast, the ‘helical-only’ Box constructs (Box A, residues 8–78; Box B, residues 94–162)^38^ only interacted with peptides corresponding to the N-terminal portion of this region (highlighted in blue, Figure S2). Notably, while previous studies have mapped the general HMGB1/CXCL12 binding surface^39–41^, our approach further refined these observations by delineating two distinct binding regions within each Box and by quantifying the contributions of specific residues. Based on these findings, we designed two additional longer constructs comprising Box A and Box B including these flanking regions (HMGB1 1-88 for Box A, HMGB1 89-174 for Box B) to assess whether inclusion of these sequences enhanced CXCL12 binding.

Biolayer interferometry (BLI, Fig. 1c, d and Figure S3) revealed that the helical-only constructs (8-78, 94-162) exhibited faster CXCL12 *k_Off_* and *k_On_* rates and weaker apparent binding affinity (higher *K_d_*) for CXCL12 compared to the two longer constructs (1-88, 89-174) which included the flanking regions. Full-length FR- and 3S-HMGB1 (1-214) exhibited similar binding kinetics to the longer constructs (Fig. 1d). Montanico et al described a 1:1 stoichiometry for FR-HMGB1–CXCL12 binding in which a single CXCL12 molecule transiently contacts residues in both HMG boxes through an intramolecular exchange mechanism^42^. Our kinetic data are in keeping with that model: although the individual HMG box cores (8–78 and 94–162) bind CXCL12 with measurable on/off rates, their interactions are relatively short-lived. Addition of the native flanking regions (1–88, 89–174) and the FL-HMGB1 substantially slow dissociation and increase apparent affinity, consistent with flanking segments stabilizing the CXCL12 contact and increasing the lifetime of the hetero-complex. These observations support a model in which CXCL12 engages a single HMGB1 molecule and samples the two Box surfaces, and where flanking regions modulate the effective kinetics and stability of that 1:1 engagement rather than establishing permanent bivalency.

Next, we investigated the importance of the flanking regions using NMR titrations with ^15^N-labeled HMGB1 constructs. As our BLI data showed equivalent binding of CXCL12 to isolated Box A and Box B as to FL-HMGB1, we performed CXCL12 titrations using the two Box B constructs we used for BLI. The extended Box B (89–174) construct displayed more pronounced chemical shift perturbations (CSPs) compared to the shorter 94–162 construct (median CSP 0.017 vs. 0.005, Figure S4a-b and Figure S5a-b), suggesting an enhanced degree of interaction with CXCL12 when the flanking regions are present.

### Identification of HMGB1 peptide regions involved in TLR-2, TLR-4 and RAGE binding

Having defined the structural determinants of the interaction between HMGB1 and CXCL12, we sought to identify regions mediating binding to proinflammatory receptors TLR-2, TLR-4/MD-2 and RAGE. We performed peptide array analysis using TLR-2, TLR-4/MD-2 and RAGE as probes to identify the specific HMGB1 sequences mediating these interactions (Fig. 2a and Figure S6). These experiments revealed that receptor binding to HMGB1 peptides is both sequence- and conformation-dependent, with oxidation status significantly influencing the structural landscape of Box A.

**Fig. 2:**
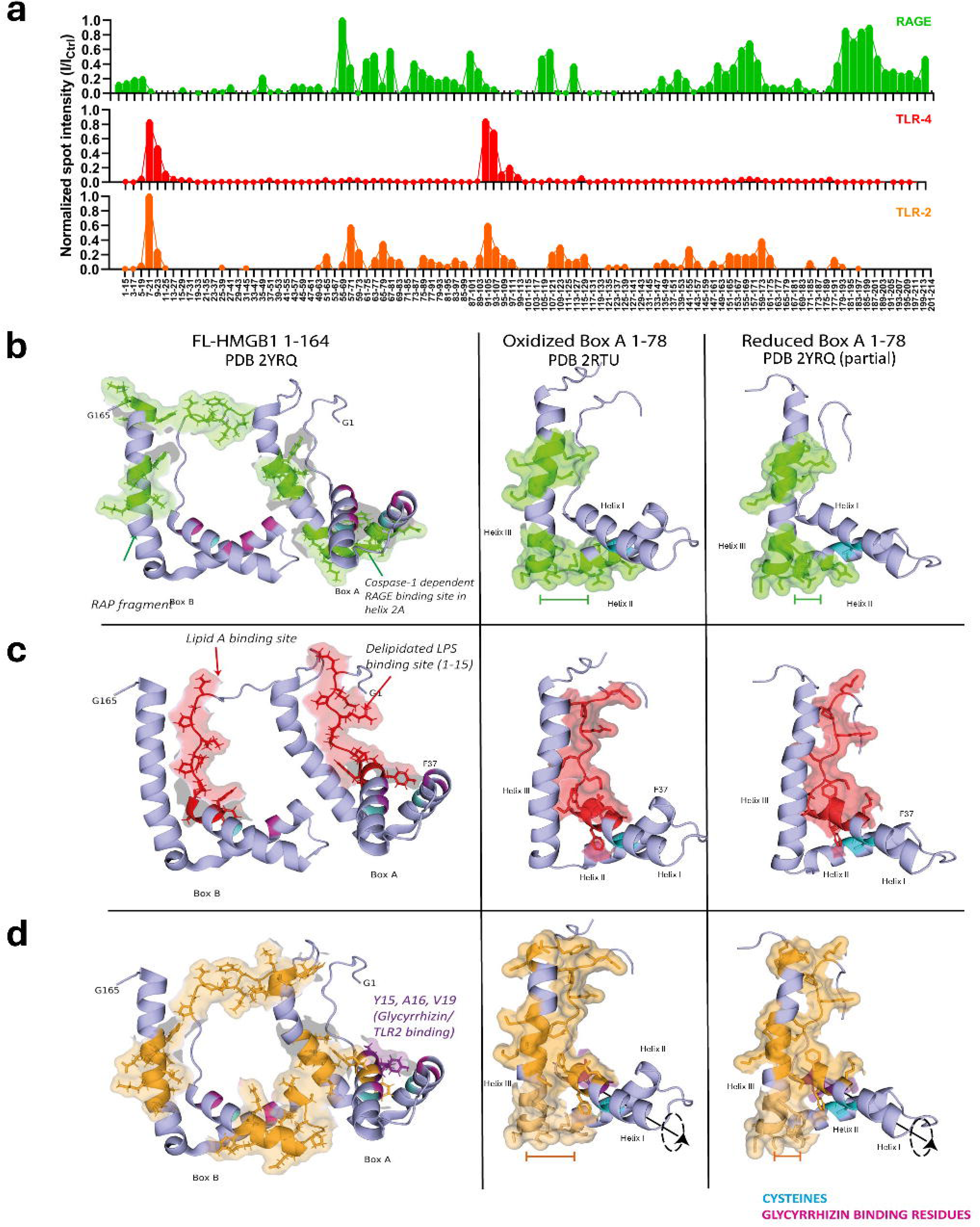
HMGB1 peptides involved in binding TLR-2, TLR-4 or RAGE reorient themselves on Box A oxidation. **(a)** Intensity densitogram (X: HMGB1 residues contained in peptide, Y: normalized spot intensity vs. control) for spot arrays of HMGB1 15-mer peptides probed with His10- RAGE (green), His10-TLR-4 (red) or His10-TLR-2 (orange) detected with anti-His HRP chemiluminescence, after subtraction of background. Baseline at 25% percentile of intensity. Surface representation of the binding peptides described overlaid on the published NMR structure of HMGB1 (PDB 2YRQ) and Box A in oxidized (PDB 2RTU) and reduced (PDB 2YRQ, truncated to only show relevant residues) states for **(b)** RAGE, **(c)** TLR-4 and **(d)** TLR-2. Cysteine residues (cyan), glycyrrhizin binding residues (pink): bars indicate distance changes between the oxidized and reduced binding regions. Axis of rotation of Helix I is also shown for TLR-2. Residues after 164 are not present in the structure and are not shown.

RAGE-binding was localized to two main regions (Fig. 2b): residues 149-182 in Box B and a second site within α-helix II of Box A, which includes the Caspase-1 activated site. These regions overlap with sequences previously implicated in RAGE modulation^43,44^. Unlike previous studies where activity was only seen following Caspase-1 cleavage of HMGB1^43^, our data show that this Box A site is solvent-accessible in full-length Box A and becomes more accessible upon oxidation due to widening of the space between helices II and III. We also observed RAGE interaction with the acidic C-terminal tail of HMGB1, a feature not shared by TLR-2 and TLR-4, suggesting a unique binding mode.

TLR-4 binding mapped to Box A, particularly within regions overlapping the LPS binding site and adjacent to the Lipid A interface^45^ (Fig. 2c). Oxidation of Box A is essential for this interaction: the presence of Cys22 within the binding interface and the redox-dependent reorientation of Phe37 in helix II^46^ (Fig. 2c) remodel the surface, creating a functionally contiguous recognition site. These oxidation-driven conformational changes provide a structural basis for the reduced ability of 3S-HMGB1 to engage TLR-4^47^. The TLR-2 interface included both α-helices I and III within Box A and partially overlapped with the glycyrrhizin-binding region^37^(Fig. 2d). Oxidation induced a redox-dependent reorientation of these helices, which likely alters the accessibility and geometry of receptor-binding surfaces involved in TLR2, as it does for TLR4, and RAGE engagement.

### Engineered dBB12L construct retains CXCL12 binding while lacking signaling via TLR-2, TLR-4 and RAGE

Next, to selectively preserve pro-regenerative signaling while eliminating pro-inflammatory receptor engagement, we engineered an HMGB1 variant where the native Box B and flanking region (residues 89–174) was retained and the N-terminal region of FR-HMGB1 (residues 1–88, encompassing Box A and its flanking residues) was replaced with a second copy of the native Box B region with its N-terminal flanking residues involved in CXCL12 binding. In this design, the native copy of Box B also contributes its C-terminal flanking residues (165–174), which are flexible and participate in CXCL12 binding, thereby ensuring that both N- and C-terminal Box B elements involved in chemokine interaction are preserved. This tandem Box B construct, designated dBB12L (FR-HMGB1 89–174::89–174) (Fig. 3a), was designed to enable interaction of each Box domain with CXCL12 in a similar manner to FR-HMGB1 while lacking critical domains for binding and signaling through TLR-2, TLR-4/MD-2 and RAGE in Box A when it is oxidized.

**Fig. 3:**
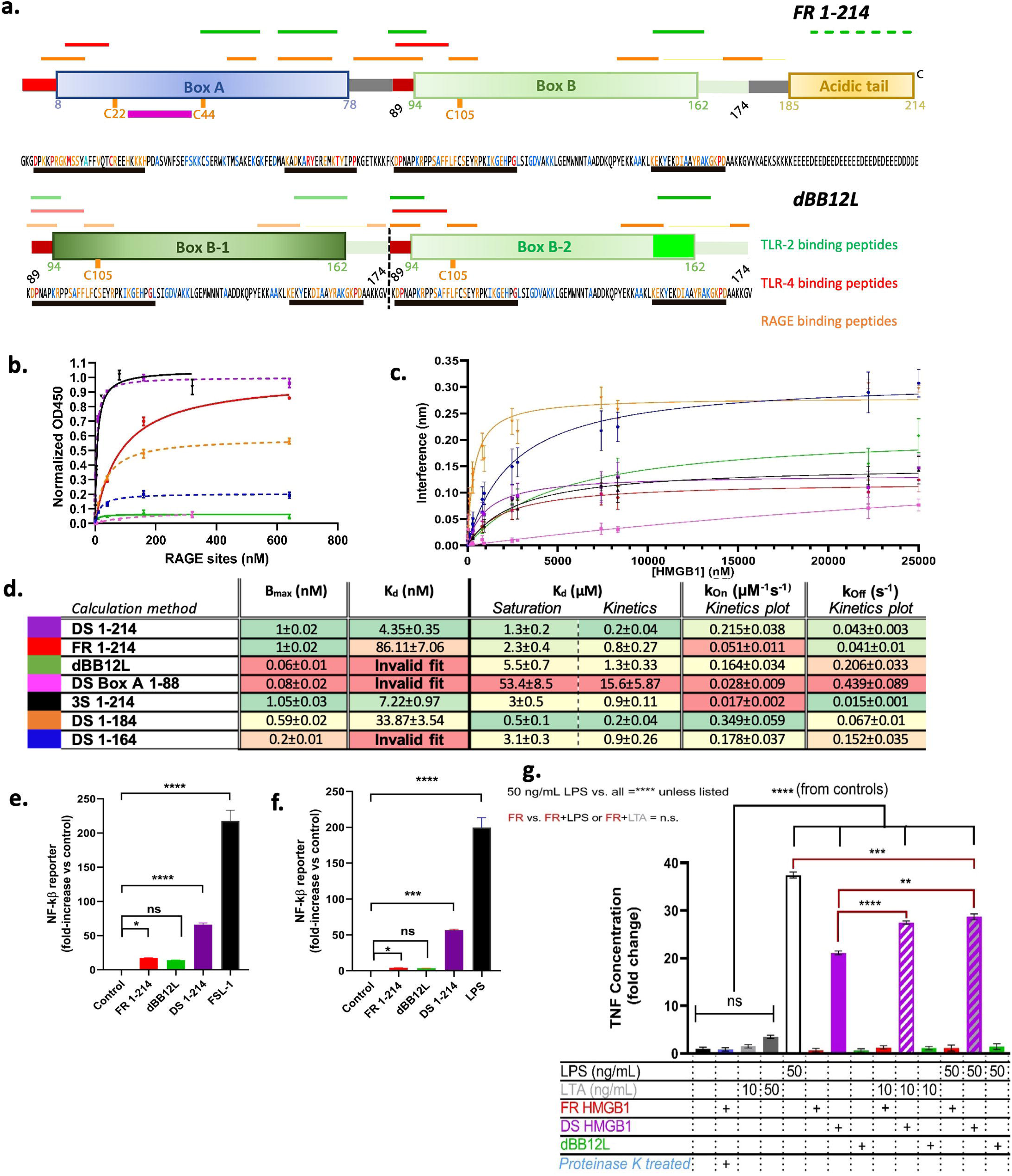
The engineered dBB12L construct that does not bind to RAGE or signal through TLR- 2 or TLR-4. **(a)** Comparison of domain organization and sequences of native FR 1-214 (top) and dBB12L (bottom). CXCL12 binding peptides are underlined below the amino acid sequences. Interaction Residues identified only by peptide arrays (red letters), identified by NMR and peptide arrays (orange letters), only identified by NMR (blue letters), are color-coded in the sequences. Horizontal bars above the domain diagrams indicate peptides binding each receptor: green (RAGE), red (TLR-4), orange (TLR-2). Vertical dashed line separates the repeat Box B units: receptor binding marks in the copy of Box B used to replace the Box A domain of FR-HMGB1 appears lighter in color. Michaelis-Menten saturation fits of HMGB1 constructs using **(b)** hybrid ELISA; n=4 per concentration, global fit normalized to DS-HMGB1 control and **(c)** BLI, 0-25 µM HMGB1; n= 6 sensor runs. In both panels, disulfide constructs are indicated by dashed lines. **(d)** Kinetic parameters are color coded in a heat map: red (weaker interaction) green (stronger interaction). B_max_ is the binding capacity (the maximum amount of a substance that can bind to all the available binding sites). Invalid fits R2 < 0.6 (poor binding). NF-κβ activity in reporter HEK-Dual cells expressing **(e)** TLR-2 and CD14 or **(f)** TLR-4, MD-2 and CD14 exposed to FR 1-214, dBB12L or DS 1-214. Data shown as mean ± SEM fold change compared to control (media alone). **(g)** TNF production by human monocytes in response to HMGB1 constructs, with or without respective co- ligands; n= 3 donors, each with 3 technical replicates.

We first assessed dBB12L binding to RAGE compared to full-length FR-, DS- and 3S-HMGB1 using solid-phase equilibrium ELISA and BLI, with additional controls included DS-HMGB1 1–184 (containing both Boxes and the RAGE-binding sequence), DS-HMGB1 1–164 (lacking the RAGE binding peptide) and DS-Box A 1–88 (negative control). By ELISA, 3S-, FR- and DS-HMGB1 showed comparable levels of RAGE binding, with DS- and 3S-HMGB1 exhibiting the highest affinities (Fig. 3b). Deletion of the RAGE-binding sequence (DS 1–164) greatly decreased binding, while deletion of the acidic tail alone (DS 1–184) had a lesser impact. Crucially, dBB12L did not bind RAGE.

Kinetic profiling of RAGE binding via BLI (Figure S7) revealed that oxidized HMGB1 constructs displayed higher association rates (*k_On_*) (Fig. 3c), while dissociation rates (*k_Off_*) were similar across constructs regardless of redox status (Fig. 3d). Truncation of the RAGE-binding region, as in dBB12L and DS 1–164, resulted in faster dissociation rates (*k_Off_*) and increased *K_d_* values, indicating weaker, more transient interactions. Among all two-Box constructs tested, dBB12L exhibited the fastest dissociation from RAGE (*k_Off_* = 0.206 s⁻¹) and the lowest binding affinity (*K_d_* = 5.5 μM, Fig. 3d), consistent with a short-lived, low-affinity complex.

We evaluated TLR-2 and TLR-4 activation by the various HMGB1 constructs in NF-κB reporter cell lines and in primary human monocytes. As expected, DS-HMGB1 activated NF-kB signaling via TLR-2 (Fig. 3e) and TLR-4 (Fig. 3f) and promoted TNF expression in primary monocytes (Fig. 3g). In contrast, neither FR-HMGB1 nor dBB12L elicited NF-κB signaling, confirming their inability to signal via these pathways. Notably, DS-HMGB1 acted synergistically with lipoteichoic acid (LTA), an effect not observed with FR-HMGB1 or dBB12L. Furthermore, when co-administered with LPS, FR-HMGB1 and dBB12L unexpectedly partially suppressed LPS-induced TNF expression, suggesting potential anti-inflammatory effects (Fig. 3g).

### dBB12L retains regenerative activity equivalent to FR-HMGB1

Having established that dBB12L is not proinflammatory, we next evaluated its capacity to promote tissue repair. In a murine skeletal muscle injury model where the tibialis anterior muscle is injected with BaCl_2_^48^, dBB12L exhibited effects equivalent to full-length FR-HMGB1 (1–214) (Fig. 4a), supporting its potential as a therapeutic to promote tissue repair. *In vitro*, using C2C12 myogenic progenitor cells, we also found that dBB12L significantly improved myogenic differentiation compared with differentiation medium (DM) alone (p < 0.001). This effect was significantly reduced in the presence of anti-CXCL12 neutralizing antibody (p < 0.01), glycyrrhizin (GL) (p < 0.05) or the CXCR4 antagonist AMD3100 (p < 0.05). These data show that, like FR-HMGB1, dBB12L-mediated muscle differentiation involves CXCL12/CXCR4 signaling (Fig. 4b).

**Fig. 4:**
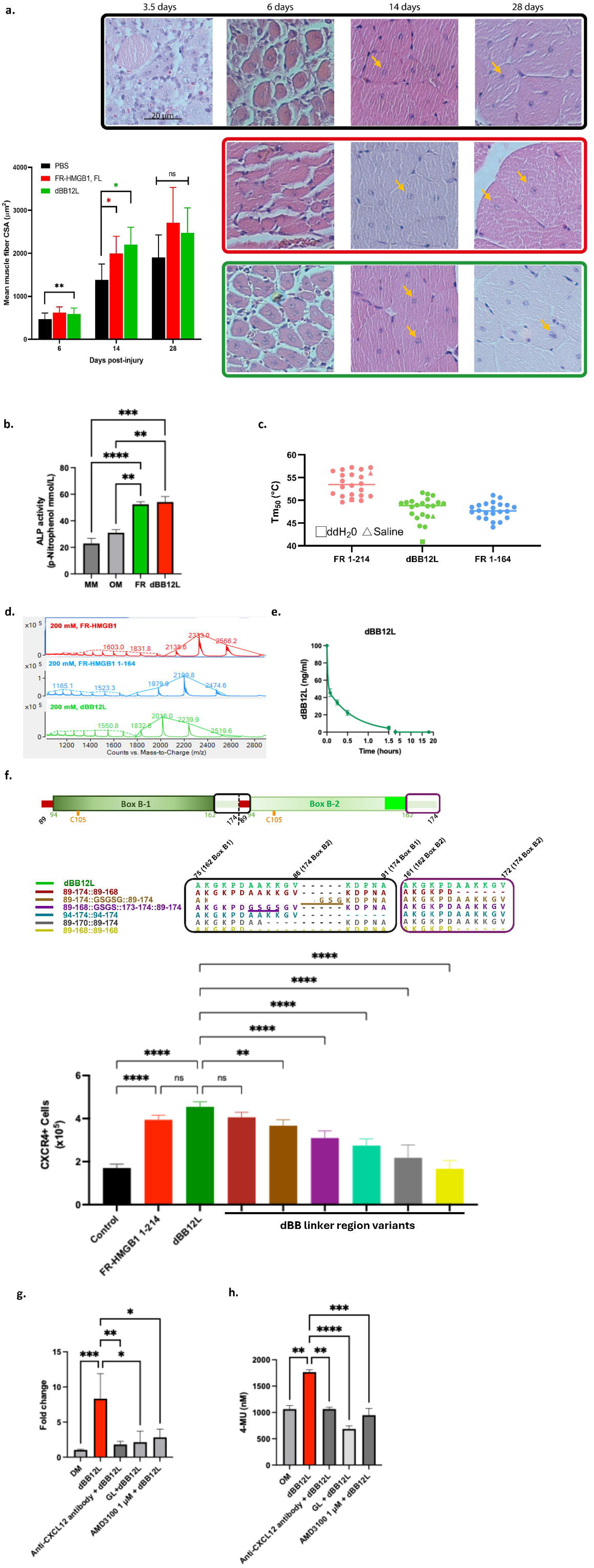
Engineered dBB12L construct retains regenerative activity similar to FR-HMGB1 and has equivalent stability and surface charge conformation to FR-HMGB1 1-164. **(a)** Mean muscle cross-section area (CSA) (95% CI) after BaCl2 injury treated with PBS (black), FR-HMGB1 1-214 (red) or dBB12L (green); n = 5 per group and time point, nested ANOVA with Holm-Sidak post-hoc correction. Representative hematoxylin and eosin images of mouse skeletal muscle treated either with FR-HMGB1 (red), dBB12L (green) or PBS control (black) after BaCl2 injury. Regenerating myofibers are identified by centrally located nuclei (indicated by arrowheads) and increased fiber diameter; both features are more prominent in FR-HMGB1- and dBB12L-treated groups compared to PBS, indicating enhanced regeneration. **(b)** C2C12 myogenic progenitor cells were induced to differentiate in the presence of dBB12L (1 µM) with or without anti-CXCL12 neutralizing antibody (10 µg/mL), glycyrrhizin (GL, 100 µM), or the CXCR4 antagonist AMD3100 (1 µM). dBB12L significantly enhanced myogenic differentiation compared with differentiation medium (DM) alone. N = 3 for each group. **(c)** Alkaline phosphatase (ALP) activity of human bone marrow derived mesenchymal stem cells treated with maintenance medium (MM), osteogenic medium (OM), dBB12L or FR-HMGB1. **(d)** Human MSCs were cultured in osteogenic medium (OM) and treated with dBB12L in the presence or absence of anti-CXCL12 antibody, GL, or AMD3100. N = 5 for each group. **(e)** Scatter plot of calculated Tm50 values for full-length, FR 1-164 and dBB12L across 24 buffer conditions. **(f)** Native ESI-MS M/Z spectra for the average folded HMGB1 monomer (ammonium acetate, 200 mM pH 6.8; theoretical effective pH under ionization 4.75 ± 1) for FR-HMGB1, FR-HMGB1(1–164), and dBB12L. **(g)** Pharmacokinetics of dBB12L in circulation following intravenous administration, fitted by nonlinear least squares to a one phase exponential decay curve, giving a half-life of 25 min. **(h)** Schematic of the tandem Box B constructs with variations on the linker length. Expression analysis of endodermal markers following HMGB1 treatment, showing dBB12L optimally promotes differentiation of endodermal progenitor cells; n = 3-5. Data are presented as mean ± SEM. Significance is indicated as *P < 0.05, P < 0.01, ***P < 0.001, ****P < 0.0001.

We have previously shown that FR-HMGB1, but not DS-HMGB1, promotes osteogenesis by human mesenchymal stem cells (MSCs)^35^, thereby highlighting the redox-dependent function of HMGB1. We also evaluated the potential of dBB12L to promote osteogenesis. Cells were cultured in maintenance medium (MM) and treated with dBB12L or FR-HMGB1 before being exposed to osteogenic medium (OM) (Fig. 4c). We found that both dBB12L and FR-HMGB1 significantly increased alkaline phosphatase (ALP) activity compared to MM (p < 0.001 for both) or OM (p < 0.01 for dBB12L; p < 0.05 for FR-HMGB1). The osteogenic effect of dBB12L was significantly attenuated in the presence of anti-CXCL12 neutralizing antibody (p < 0.01), glycyrrhizin (GL) (p < 0.0001) or the CXCR4 antagonist AMD3100 (p < 0.001), indicating that the pro-osteogenic activity of dBB12L is at also dependent on CXCR4 signaling and is attenuated by disruption of the HMGB1–CXCL12 interaction (Fig. 4d).

If dBB12L is to be developed as a therapeutic, it would be important to assess its stability. Therefore, we assessed the biophysical characteristics of dBB12L (Box B–linker–Box B) relative to FR-HMGB1 1–164 (Box A–linker–Box B) by comparing their stability and conformational properties. Both constructs displayed similar thermostability by differential scanning fluorimetry (DSF) (Fig. 4e) and comparable surface charge distributions and conformational states by native ESI/MS (Fig. 4f). Both constructs existed as monodisperse populations with two major charge states: a compact monomeric form (higher m/z) and a more relaxed conformation (lower m/z), indicating similar folding and solution behavior. Both variants remained monodisperse despite displaying multiple charge state species and were stable in solution for at least 180 days without preservatives (Figure S8). Pharmacokinetic analysis revealed that dBB12L had a circulating half-life of approximately 24.6 minutes following intravenous administration in mice (Fig. 4g), compared to 24 min for FR-HMGB1 1–164^35,49^.

Next, we generated more tandem Box B variants by changing the dBB12L linker length and composition and probed their influence on the regenerative activity by measuring their ability to induce differentiation of CXCR4⁺ endodermal progenitors (Fig. 4h). Deletion of the N-terminal linker region (residues 89-93 in wild type HMGB1: KPDNA) in the first Box B (yielding FR-HMGB1 94-174::94-174) reduced activity, consistent with the lower CXCL12 binding affinity for Box B 94-162 observed using BLI. Whilst deletion of residues 169-174 from the C-terminus of the second Box B in dBB12L did not affect activity (FR-HMGB1 89-174::89-168), deletion of residues 171-174 from the first Box B (shortening the linker region to 13 residues (FR-HMGB1 89-170::89-174) reduced activity, despite retention of CXCL12 binding residues. Complete deletion of the C-terminal regions from both HMG Box B domains that do not bind CXCL12 (FR HMGB1 89-168::89-168) almost completely abolished activity. Insertion of a Gly-Ser spacer after the first Box B (FR HMGB1 89-174::GSGSG::89-174) or replacing non-CXCL12 binding residues with Gly-Ser spacers in the same region (FR HMGB1 89-168::GSGS::173-174::89-174) both led to reduced activity. These results indicate that a linker length of approximately 13–22 amino acids between the two Box B domains is optimal for promoting repair, and that deviations in either direction compromise regenerative activity.

## Discussion

Our study provides a detailed structural and functional map of HMGB1 interactions with CXCL12 and key pro-inflammatory receptors, revealing how domain architecture and redox state influence biological outcomes. Systematic mapping of HMGB1–CXCL12 interactions revealed that both Box A and Box B domains independently bind CXCL12 via homologous regions across HMG Boxes located on the underside of each domain, and the flanking regions surrounding the canonical HMG Boxes significantly stabilize this interaction. These findings refine the current model of the HMGB1-CXCL12 interface by demonstrating that CXCL12 engagement extends beyond the minimal structured HMG domains^41^ and involves distributed interaction motifs that modulate complex stability.

Although CSPs are small (with the largest CSPs on the 0.1-0.2 ppm range) the magnitudes observed fell within the range reported for full-length HMGB1 in previous NMR studies^42^. Notably, these analyses were performed under near-physiological buffer conditions (150 mM NaCl, pH 7.4). BLI revealed comparable affinities of the individual Box domains for CXCL12, and similar binding regions were also observed in the peptide arrays. These data suggest that the two Box domains differ primarily in binding geometry or interaction dynamics rather than equilibrium affinity. Inclusion of flanking residues reduced CXCL12 dissociation rates without markedly altering association kinetics, indicating that these regions contribute to stabilization of chemokine presentation. Collectively, these observations support the dynamic model in which CXCL12 can associate with either Box HMG domain and potentially transfer between them^42^.

Importantly, our data clarify distinct functional roles for the two HMG domains. While both Box A and Box B can bind CXCL12, inflammatory receptor engagement was strongly dependent on oxidation state and domain context. Oxidized Box A mediated robust binding to TLR-2, TLR-4 and RAGE, with peptide mapping indicating that TLR-4 recognition involves a discontinuous surface in Box A that becomes functionally contiguous upon oxidation. For RAGE, oxidized Box A appears to facilitate an initial interaction that is further stabilized by contributions from the C-terminal region. These data support a biphasic model of RAGE interaction^50^: oxidized Box A first mediates the initial, oxidation-dependent low-affinity contact, which is then stabilized by high-affinity binding via the C-terminal region. TLR-2 binding was largely confined to glycyrrhizin-sensitive regions within the core Box domains^12,37,51^, with oxidation of Box A facilitating these interactions, whereas the acidic tail did not appear to contribute significantly to TLR-2 engagement. In contrast, Box B displayed limited capacity to engage these inflammatory receptors in our assays while retaining CXCL12-binding activity. This functional divergence suggests that Box A acts as a redox-sensitive determinant of inflammatory signaling, whereas Box B provides a structurally stable module capable of supporting CXCL12-mediated regenerative signaling.

Leveraging these mechanistic insights, we engineered tandem Box B construct dBB12L to preserve CXCL12 engagement while excluding the oxidation-sensitive Box A domain and the acidic tail. A previous study proposed that the negatively charged acidic tail facilitates CXCL12 recruitment through long-range electrostatic interactions, thereby promoting subsequent engagement of Box A^42^. Consistent with this model, deletion of the acidic tail altered CXCL12-binding properties in truncated FR-HMGB1 constructs. However, despite lacking both Box A and the full acidic tail, dBB12L retained regenerative efficacy *in vitro* and *in vivo* in the models tested. Importantly, its effects were abolished by CXCL12 neutralization and blockade of CXCR4, confirming its dependence on the CXCL12–CXCR4 axis. These data indicate that efficient CXCL12-mediated regenerative signaling does not strictly require Box A or the acidic tail, but rather sufficient CXCL12 engagement and appropriate spatial presentation of two Box domains. Duplication of Box B, combined with removal of potential tail-mediated competition, may compensate for differences in domain-specific interactions, enabling formation of a functional heterocomplex that effectively signals via to CXCR4 to promote tissue repair. At the same time. the dBB12L construct exhibited markedly reduced binding to RAGE and signaling via TLR-2 and TLR-4. dBB12L did not activate NF-κB or promote TNF expression, in contrast to DS-HMGB1, and even suppressed LPS-induced TNF production, suggesting a partial antagonist effect. It also failed to synergize with LTA in TLR-2 activation assays, supporting canonical inflammatory receptor activation can be functionally separated from regenerative CXCL12 signaling through structural redesign of HMGB1.

Beyond its functional selectivity, dBB12L retained favorable biophysical characteristics, including thermostability, solubility and prolonged solution stability. Native mass spectrometry and size exclusion chromatography confirmed that the construct remains monomeric and conformationally dynamic in solution, resembling reduced HMGB1 while lacking detectable oxidation-dependent inflammatory receptor engagement. These properties support its potential suitability for further therapeutic development.

Our study further demonstrates that the spatial geometry between the two Box B domains plays a critical role in maintaining regenerative activity. Regenerative activity was sensitive to linker length: shortening the linker impaired activity, particularly when CXCL12-stabilizing flanking residues were removed, whereas excessive extension also reduced efficacy. Within the range tested, a linker length of approximately 13–22 residues appeared to provide an appropriate balance between flexibility and spatial proximity. These results are consistent with recent findings where the HMGB1-CXCL12 interaction is highly dynamic and does not form a static complex^42^. Previous studies showed that CXCL12 binding is preserved across individual Box domains of full-length FR-HMGB1, with no evidence of cooperative binding, indicating that each Box binds monomeric CXCL12 independently^52^. Our observations also support the dynamic model in which CXCL12 can transfer between HMG domains^45,42^, with Box A exhibiting slower dissociation kinetics than Box B. Excessive distance between HMG Boxes may impair the exchange of CXCL12 between the two Box domains, and a linker region shorter than 13 amino acids would likely hinder the conformational changes needed for the heterocomplex to bind CXCR4 or to support interchange of CXCL12 between HMG Boxes.

In summary, our findings provide a framework for designing HMGB1-based constructs that promote regeneration without the potential for deleterious inflammatory or pro-thrombotic effects and supports a modular view of HMGB1 in which its regenerative and inflammatory functions can be structurally and functionally separated. By preserving CXCL12-binding motifs while removing oxidation-sensitive and C-terminal acidic tail-mediated receptor interactions, dBB12L emerges as a promising therapeutic for promoting tissue repair without the risk of triggering potentially deleterious inflammation or thrombosis.

## Limitations

While our study demonstrates that the engineered tandem Box B construct dBB12L effectively decouples tissue regeneration from pro-inflammatory signaling, several limitations warrant consideration. First, our evaluation of RAGE engagement by dBB12L was limited to *in vitro* biochemical binding assays (ELISA and BLI). The lack of an established cell-based reporter for RAGE without cross-interference from other NF-κB-activating receptors precluded the direct measurement of downstream RAGE signaling in cell culture. While the complete absence of detectable RAGE binding strongly supports our conclusion that dBB12L does not act through this axis, the future development of highly specific RAGE reporter systems will be necessary to definitively confirm the absence of downstream signaling in complex cellular environments.

Our *in vivo* and *in vitro* models were based on restricted evaluation of sex as a biological variable. Specifically, the acute skeletal muscle injury model utilized exclusively female C57BL/6J mice and the primary human CD14+ monocytes used for inflammatory assays were sourced solely from male donors. Consequently, potential sex-dependent dimorphisms in receptor expression, baseline inflammatory responses, or regenerative kinetics remain unaddressed, limiting the immediate generalizability of these findings.

Finally, although we validated the regenerative efficacy of dBB12L in a localized acute injury model, its performance in complex tissue environments characterized by severe chronic ischemia or extensive fibrosis remains unexplored. For example, native FR-HMGB1 supports functional recovery following myocardial infarction^36^. Hence evaluating dBB12L in a similar model will be an important next step.

## STAR METHODS

### Key Resource Table

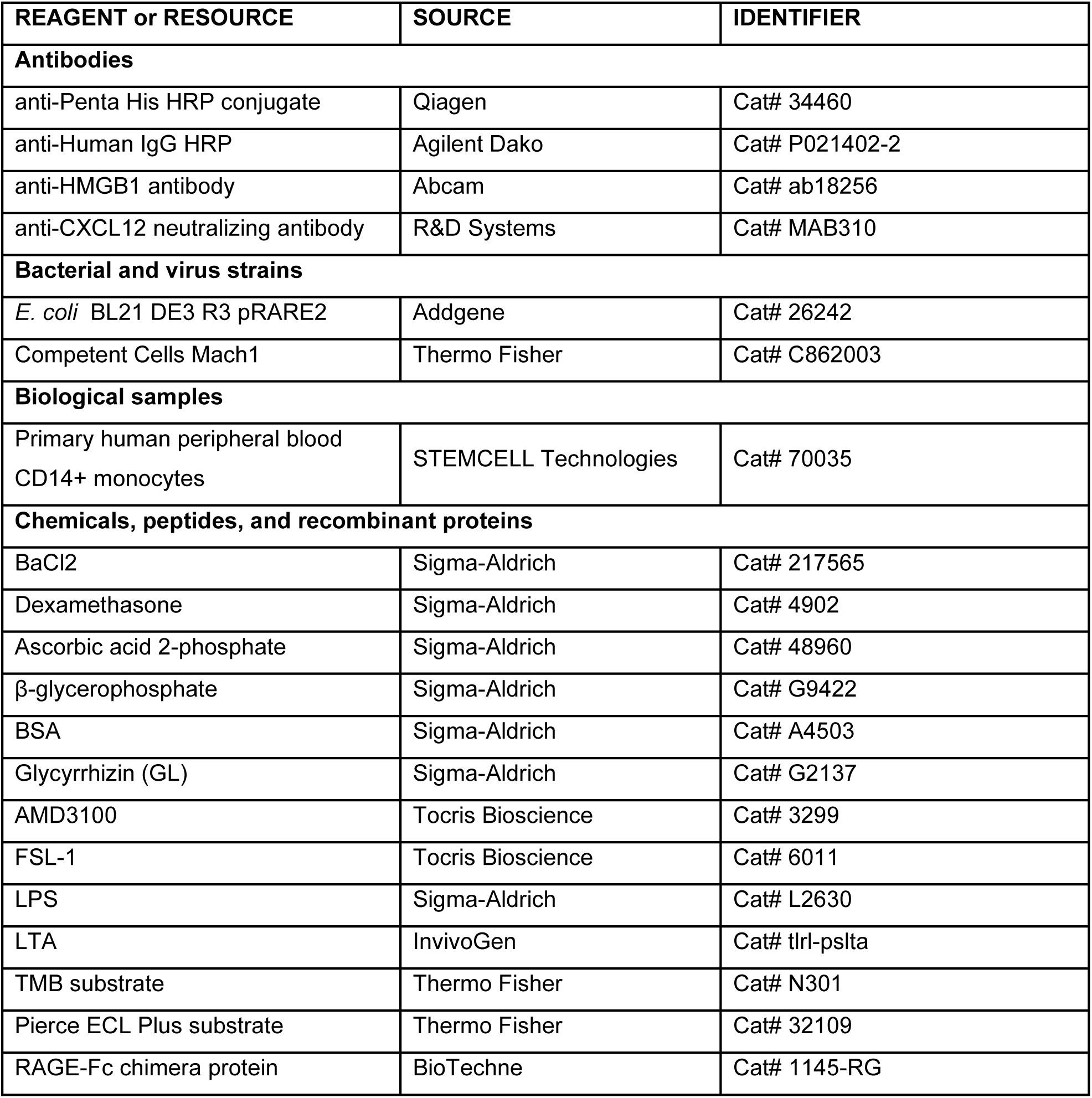
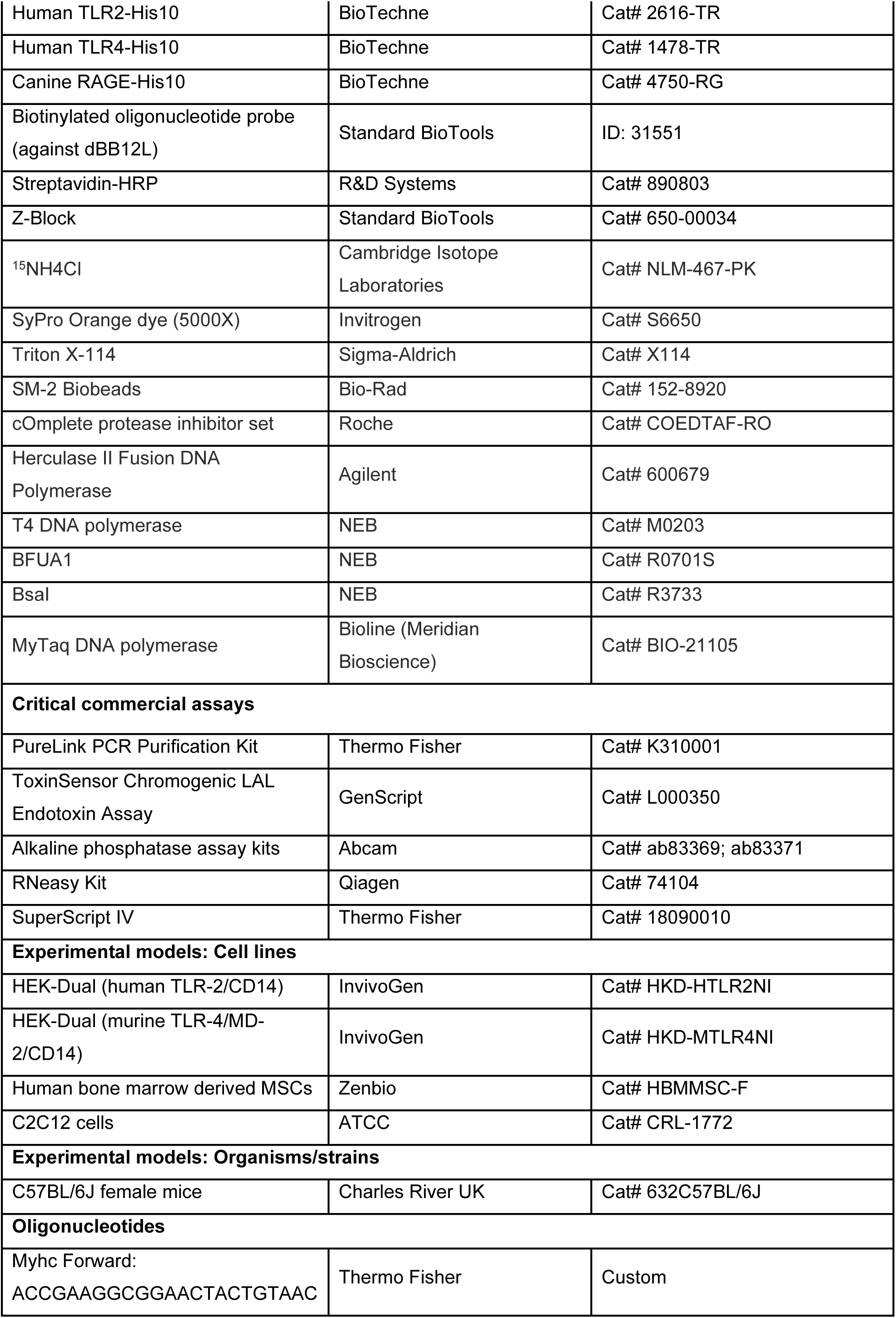
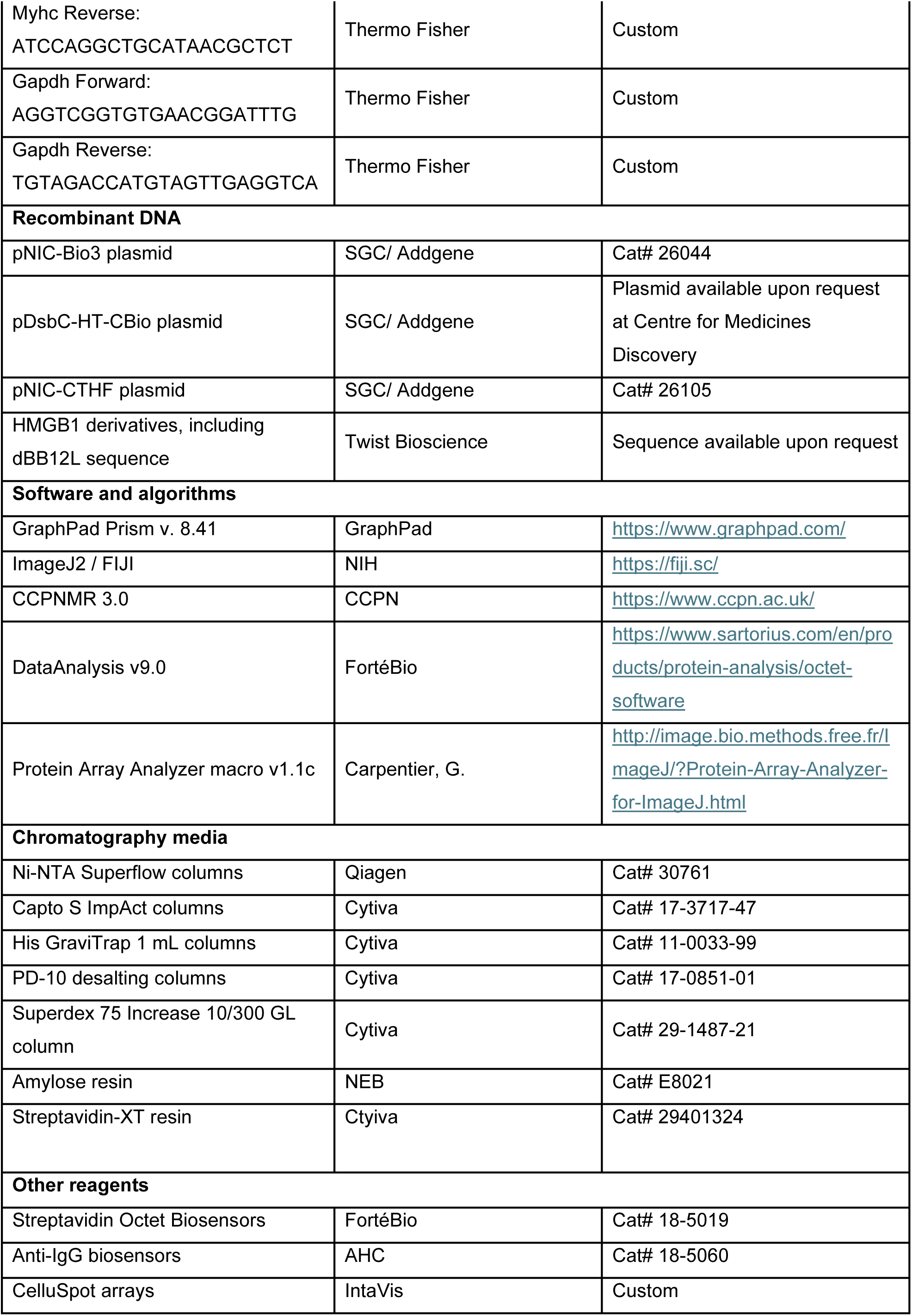

## RESOURCE AVAILABILITY

### Lead contact

Further information and requests for resources and reagents should be directed to the corresponding author, Prof. Jagdeep Nanchahal.

## Materials availability

Requests for plasmids, DNA maps and engineered proteins should be directed to the corresponding author.

## Data and code availability

All data reported in this paper will be shared by the corresponding author upon request. This paper does not report original code. Any additional information required to re-analyze the data reported in this paper is available from the corresponding author upon request.

## EXPERIMENTAL MODEL AND STUDY PARTICIPANT DETAILS

### Animal models

C57BL/6J female mice aged 10-11 weeks were purchased from Charles River UK and allowed to acclimatize for 1 week. All animal procedures were approved by the University of Oxford Animal Care and Ethical Review Committee and the United Kingdom Home Office (PPL 30/3330 and PPL P12F5C2AF)). Only female mice and only male human monocyte donors were utilized. Therefore, the influence of sex on these results cannot be determined, which represents a limitation of this study’s generalizability.

### Primary cell cultures

Primary human peripheral blood CD14+ monocytes (Catalog #70035) were purchased from STEMCELL Technologies. The cells were sourced from three healthy, non-smoking Caucasian male donors (Donor IDs: 110042888 [Age 30], 110039568 [Age 34], and 110042930 [Age 30]). Monocytes were maintained in DMEM (Gibco), supplemented with 10% FBS (Gibco) in standard tissue culture conditions (37°C; 5% CO2).

Human bone marrow derived mesenchymal stem cells (hMSCs, Zenbio) were maintained in DMEM (Gibco) supplemented with 10% FBS (Gibco), 1% L-glutamine (GE Healthcare) and 1% penicillin/streptomycin (GE Healthcare) in standard tissue-culture conditions (37 °C; 5% CO2) and used between passages 3–5

### Cell lines

HEK-Dual cells (InvivoGen) expressing human TLR-2 and CD14 or murine TLR-4, MD-2 and CD14 were maintained in DMEM (Gibco), supplemented with 10% FBS (Gibco), 1% L-Glutamine (Gibco) and 1% penicillin/streptomycin (Gibco), in standard tissue culture conditions (37°C; 5% CO2). C2C12 cells (ATCC) were cultured in DMEM (Gibco) supplemented with 10% FBS (Gibco), 1% L-glutamine (GE Healthcare), and 1% penicillin/streptomycin (GE Healthcare) under standard tissue culture conditions (37 °C, 5% CO2)

## METHOD DETAILS

### Bacterial culture media

SOC: 20 g/L tryptone, 5 g/L yeast extract, 0.5 g/L NaCl, 0.1862 g/L KCl were autoclaved and supplemented with 4.132 g/L MgCl2 and 20 mM glucose. LB (Luria Bertani): 10 g/L tryptone, 5 g/L yeast extract, 10 g/L NaCl, pH 7.2, autoclave sterilized. 2% w/v agar powder was added to make LB agar plates. TB (Terrific Broth): 12 g/L tryptone, 24 g/L yeast extract, 0.4% glycerol, 12.5 g/L K2HPO4, 2.35 g/L KH2PO4, autoclave sterilized. TB supplemented: TB formula plus 1.6% w/v glycerol, 10 g/L glucose, 25 mM (NH4)2SO4, 10 mM MgSO4, 10X trace metals, 0.22 µM sterile filtered. Trace metal solution: 50 mM FeCl3 (13.5 g/L), 20 mM CaCl2 (2.94 g/L), 10 mM MnCl2 (1.96 g/L), 10 mM ZnSO4 (2.88 g/L), 2 mM CoCl2 (0.48 g/L), 2 mM CuCl2 (0.34 g/L), and 2 mM NiCl2 (0.48 g/L), in 0.1 M HCl, 0.22 µM sterile filtered. M9 minimal medium: 16 g/L Na2HPO4, 4 g/L K2HPO4, 1 g/L NaCl, pH 7.2-7.3 and 2.5 g/L FeSO4, 0.25 mg/L ZnCl2, 0.05 mg/L CuSO4, 0.25 g/L EDTA, 1 mM MgSO4 were autoclaved and supplemented with 4 g/L glucose, 1 g/L 15NH4Cl, 0.3 mM CaCl2, 1.5 mg/L D-biotin and 1.5 mg/L Thiamine-HCL from sterile filtered stocks.

### *E. coli* strains

Chemically competent stocks of Mach-1 T1R, BL21(DE3)-R3-pRARE2 (chloramphenicol resistance 36 µg/mL, T7-polymerase *lac* induction57), and BL21(DE3)-R3-pRARE2-BirA (additional spectinomycin resistance 50 µg/mL for *in vivo* biotinylation) were used for plasmid storage and recombinant protein expression.

### Plasmids

All expression plasmids utilized contained a 6xHis tag with a TEV-cleavage site. The pNIC-Bio3 and pDsbC-HT-CBio vectors incorporated C-terminal biotinylation epitopes (which can be removed with a stop codon). Plasmid DNA was linearized by restriction enzyme digestion: BfuA1 (3 h, 60°C) for pNIC-CTHF or BsaI (2 h, 37°C) for pDsbC-HT-CBio and pNIC-Bio3 Cut vector DNA was purified and treated with T4 DNA polymerase in the presence of 0.25 mM dGTP (pNIC-CTHF) or dCTP as per manufacturer protocols.

### Cloning

HMGB1 constructs sourced from the Mammalian Gene Collection were amplified via PCR: a program of 95°C/10’, 25x (95°C/30”, 52°C/1’, 0.5-1.5’ at 68°C), 68°C/10’ was used. Reaction consisted of 5 µL Herculase II buffer, 1 µM of each primer, 6 µg/mL plasmid template, 1 µM dNTP mixture and 1 unit Herculase II Fusion DNA Polymerase in 25 µL final volume. PCR products were purified using a PureLink PCR Purification Kit before further use. Amplified coding sequences (alleles) were cloned into the destination vector via ligation independent cloning (LIC). The insert was treated with T4 DNA polymerase in the presence of a cognate nucleotide to that used for the vector (10 µL reaction volume), and 2 µL was mixed with 1 µL of treated vector and annealed for 30‘. 40 µL ice-cold Mach-1 cells (for storage) or 20 µL BL21(DE3)-R3-pRARE2/BL21(DE3)-R3-pRARE2-BirA cells (for expression) were added and heat-shocked for 45” at 42°C before chilling on ice. Recovery was performed for 2 h in SOC medium at 37°C prior to plating on selective media with 5% sucrose and antibiotics. After 24 h, positive colonies were picked and screened with MyTaq DNA polymerase according to manufacturer protocols with specific sequencing primer pairs for bands of the correct molecular weight. Positive transformants were grown overnight in 1 mL of 2X LB (double concentration of LB) with antibiotics and stocked with 12% glycerol v/v at -80°C.

3S-HMGB1 mutant sequence was generated in a similar manner. PCR was performed separately to generate fragments containing the C22S/C44S and C105S mutations, which were subsequently annealed via PCR; 5 µL of each purified PCR product substituted the primers and template in this reaction. CXCL12 constructs were cloned with an in-frame SUMO protease site N-terminal to the mature protein to allow for periplasmic secretion with an N-terminal fusion protein in the pDsbC-HT-CBio vector (DsbC-SUMO-CXCL12) to avoid addition of N-terminal residues to the protein which could affect its activity^53,54^ whilst obtaining folded, oxidized CXCL12 via the DsbC fusion protein system^55^. All mutants in this work were verified by sequencing (Source Biosciences). The sequence for HMGB1-dBB was designed *in silico* by codon-optimizing a Box B 89-174 sequence according to *E. coli* BL21-DE3 genome (assembly ASM956v1) placed after the native HMGB1 Box B sequence, and synthetized *in vitro* and cloned in pNIC-CTHF. Plasmid stocks were stored in Mach 1 cells.

### Recombinant protein expression

20 mL of overnight culture of HMGB1-expression strain transformants, grown from a fresh agar plate streak, were inoculated into 1 L of TB (or M9) medium with supplement and allowed to grow up to OD 2.0 at 37°C with 0.45 RCF orbital shaking (OD 0.6 for M9 medium). Precultures used for production of ^15^N labelled HMGB1 were first spun down at 1000 RCF for 5 ‘and washed in M9 medium. Once the target OD was reached, were cooled to 18°C before addition of 0.5 mM or 0.25 mM IPTG (for HMGB1 and CXCL12 proteins respectively) and grown for 16 h before harvesting at 4000 RCF. For biotinylated proteins, 10 mM D-biotin in PBS was added before induction and again 1 h before cell harvesting.

### Recombinant HMGB1 purification

Pellets of induced HMGB1-expressing cells were resuspended at 14 g/L in 1 M NaCl, 5% glycerol, 50 mM HEPES pH 7.5, 10 mM Imidazole (Buffer A) supplemented with 1:1000 protease inhibitors, 3 µg/mL Benzonase-MBP, 1 mM MgSO_4_, 0.5 mg/L lysozyme and 0.5% v/v Triton-X100 before freezing at -80°C; from this point onwards all steps took place at 4°C. Thawed pellets were spun down at 6780 RCF for 45’ and the supernatant was loaded into pre-equilibrated His GraviTrap 1 mL columns. After drip-through, columns were washed with 10 CV of 1 M NaCl, 50 mM HEPES pH 7.5 and 1.5 CV of 0.4 M NaCl, 20 mM HEPES pH 7.5, 1 mM MgSO_4_, and 3 µg/mL Benzonase-MBP solution to digest remaining DNA for 30‘. Contaminants were washed with 15 CV of 0.5 M NaCl, 5% glycerol, 50 mM HEPES pH 7.5 (Buffer B) supplemented with 30 mM imidazole before elution directly into a PD-10 desalting column (equilibrated in Buffer B + 20 mM imidazole) with 2.5 mL of Buffer B + 500 mM imidazole. Proteins were eluted from the column with 3.5 mL of Buffer B + 20 mM imidazole before tag removal with 1:20 OD TEV-GST protease over 16 h.

Proteases and further contaminants were removed by recirculating the protein solutions over the same GraviTrap column used to purify initially (equilibrated in Buffer B+20 mM imidazole). For biotinylated proteins, Streptavidin-XT resin was used instead to select biotinylated molecules only: after 30‘ of incubation in the resin, sample was allowed to drip through, washed with 30 CV of buffer A and 1 CV of buffer B + 100 mM D-biotin, and eluted by incubation in 3 CV of the same buffer for 2 h. Proteins were further purified by size exclusion chromatography (SEC) (Superdex 75 Increase 10/300 GL column at 0.35 mL/min or 16/600 at 1.2 mL/min flow rate) in either 10 mM HEPES pH 7.5 + 150 mM NaCl for biophysics work or cell-culture grade PBS for cell and animal work. Recombinant proteins were flash-frozen in liquid nitrogen for storage, adding 1 mM TCEP in the case of reduced HMGB1 proteins.

### Recombinant CXCL12 purification

Outer membranes of cells expressing DsbC-SUMO-CXCL12 were lysed by osmotic shock^56^. Pellets were resuspended at 40 g/L in 1 M sucrose, 0.2 M Tris-HCl pH 8.0, 1 mM EDTA, 1 mg/mL lysozyme, 2X cOmplete protease inhibitor set, 50 mM Imidazole and 3 µg/mL benzonase. This was stirred for 45 minutes at room temperature before adding 4 volumes of ice-cold 18.2 mΩ water and mixed for a further 10 minutes before adding 1 mM MgSO_4_. This was centrifuged for 1 h at 16000 RCF, 4°C, and the supernatant loaded at 10 mL/min into Ni-NTA Superflow columns on an Äkta Xpress FPLC system; 1 column was used for every 6 L of cells. Proteins were eluted via an imidazole gradient (10-25 mM over 10 CV, and 25-500 mM over 8 CV) in Buffer B, and 1:10 OD of Ulp-1 protease were added before dialysis in 100 volumes of 0.2 M NaCl, 20 mM HEPES pH 8.0 (Buffer Ac) overnight. On the next day, the protein was loaded into CaptoS ImpAct columns and eluted in a gradient of 0.2-1.5 M NaCl in 20 mM HEPES pH 8.0 to separate cut CXCL12 from DsbC and Ulp-1. Proteins were further purified via SEC in the same way as HMGB1 and flash-frozen for storage. HEPES buffers were used instead of phosphate-based formulations to prevent CXCL12 dimerization in any downstream assays, therefore focusing only on the monomeric form of the protein^57^. Correct oxidative state of CXCL12 was verified by reverse-phase HPLC as published^58^.

### Removal of endotoxins

Endotoxin was removed in all cases before size exclusion chromatography via phase separation with Triton X-114^59^. A 2% v/v of TX-114 was added to recombinant protein solutions, homogenized for 20’ with orbital shaking at 2000 RCF at 4°C, and separated for 5’ at 37°C before pelleting the detergent phase at 8000 RCF, 10’, 25°C. The supernatant was mixed with 5% w/v of SM-2 Biobeads, cleaned with 2% TX-114 for 2 h and regenerated with 30 CV of methanol, 30 CV of endotoxin-free 18.2 mΩ water and 30 CV of endotoxin-free PBS. This was incubated for 4 h at room temperature to adsorb remaining Triton and PEG^60^ before injection onto a sanitized SEC system (with 0.5 M NaOH contact over 12 h, followed by 0.2 M acetic acid/20% ethanol contact over 6 h and equilibration in cell-culture grade PBS) to fully remove leftover polymer contaminants whilst performing size exclusion. The absence of Triton and PEG was verified by lack of their respective charge state species in ESI/QTOF-MS mass spectrometry: due to the ion-suppressant nature of both compounds, any trace of either would immediately overwhelm the protein signal during ionization^61^. LPS content of the recombinant proteins was assessed via the ToxinSensor Chromogenic LAL Endotoxin Assay. Samples were approved for cell and animal use when they contained < 1 EU LPS/mg protein.

### Enzyme production

TEV-GST protease (GST-fusion protein), Benzonase-MBP, and Ulp-1 protease were produced from transformants in storage at the SGC collection^62^; all had 200 µg/mL ampicillin resistance. TEV and Ulp-1 were purified as per the protocols described for HMGB1 with only one IMAC step, whereas Benzonase-MBP was purified from outer membrane lysates obtained as with CXCL12 and isolated with use of amylose resin as per manufacturer protocols. In both cases, the resulting proteins were concentrated to 10 mg/mL in 50 mM HEPES pH 7.5, 0.3 M NaCl, 10% glycerol. GST-TEV protease and Ulp-1 were flash-frozen with liquid nitrogen and supplemented with 0.5 mM TCEP during purification; Benzonase-MBP was supplemented with 50% glycerol and 2 mM MgCl_2_ and stored at -20°C.

### Differential scanning fluorimetry (DSF)

Thermal transition temperature (Tm50) was measured via emission of fluorescence in the presence of SyPro Orange dye. 24 μL of different reduced HMGB1 constructs at 0.5 mg/mL (4 μM) in 10 mM HEPES, 150 mM NaCl, pH 7.5, 1 mM TCEP were premixed with SyPro orange dye at a final concentration of 25X and added to a final volume of 120 μL of liquid in the respective buffers (Table S1), on a white qPCR fluorescence plate and sealed with a fluorescence-compatible seal. Thermal denaturation was induced by preincubating the plate at room temperature for 15 minutes, then heat at 1°C intervals between 25 and 85°C. Measurements were taken 1 minute after each temperature increase in a StrataGene MXPro3500p instrument. For each protein, three wells were taken in replicate and fitted individually. Change in fluorescence was extracted as R (multicomponent view) and fitted to a Boltzmann sigmoidal curve to extrapolate Tm50. Fitted values were averaged across each buffer condition/protein pair.

### Mass spectrometry

For protein identification via MS/MS and tryptic digest, bands from SDS-PAGE gels were excised and submitted to the open access MS platform and analyzed by Dr. Rod Chalk, Tiago Moreira and Oktawia Borkowska, as published^62,63^. Data analysis (peptide mapping) to annotate protein identity was performed with the MASCOT search engine, against the Uniprot and SGC databases. Native ESI/MS experiments were performed by manual injection in volatile buffer (50 or 200 mM ammonium acetate, pH 6.5) into an ESI/QTOF instrument at 360 µL/h. Signal was acquired for at least 10 counts (30 sec) once a steady ionic stream was observed in the total ion chromatogram. For denaturing experiments, samples were diluted to 1 mg/mL in 0.2% formic acid and injected via HPLC and eluted in a mobile phase of formic acid/methanol, as described65; lower mW species were identified via the use of the PAWS peptide mapping software. Each continuous distribution of charge states was considered a distinct conformation; charge states (Z) were assigned according to the formula where mW= (mW/Z-proton mass)* Z. Surface areas were derived from the formulas proposed in the literature^64,65^. At least three independent injections were performed for all MS samples.

### Analytic SEC experiments

Standard protein sets of known molecular weights were run through a Superdex 75 Increase 10/300 GL column used in these experiments; injections were performed at 1 mg/mL to avoid saturation of signal and all samples were eluted at 0.4 mL/min. Values shown are all from triplicate experiments. Between runs, the column was re-equilibrated in reverse flow under the same conditions with 2 CV buffer.

### Peptide arrays

In-house peptide arrays were used for the screening of HMGB1 (Uniprot P09429) and CXCL12 (Uniprot P48061) sequences, printing peptides containing 15-mer overlapping sequences of each protein shifted by 2 amino acids each time. For alanine scanning of HMGB1 hits on the initial array, membranes with point mutations to alanine of the identified peptides were printed by the same method. Residues whose mutation to alanine resulted in higher intensity changes than those observed for alanine positions in the sequence were considered as significant contributors to CXCL12 binding.

For probing interactions with DAMP receptors, CelluSpot arrays (IntaVis) were used due to cost considerations. Corrected values were normalized again from 1 to 0 for comparability across datasets. An additional HMGB1 peptide array was synthesized on the same support to confirm consistency across the different membrane types (data not shown). Full sequences of printed peptides in arrays are present in Supplemental Information as an .xslx file; for the Alanine scan, sequences printed are shown in the relevant figures.

For both types of arrays, membranes were rehydrated with 95% and 70% ethanol at 20-25°C, equilibrated with PBST (PBS 1X + 0.05% Tween-20, 3x), and blocked with 10% BSA/PBST for 8 h. 1 µM of the partner His-tagged protein construct (in PBS) was added and allowed to bind for 24 h at 4°C. For DAMP receptor assays, human TLR2-His10, human TLR4-His10, or canine RAGE-His10 (all from BioTechne) were used; HMGB1 and CXCL12 proteins were in-house purified. Excess BSA and protein were removed with 3 washes in PBST; all washes lasted 1 min unless otherwise stated. Bound proteins were detected by treating the membranes with 1:3000 dilution of Qiagen anti-Penta His HRP conjugate and excess antibody removed with 3 washes in PBST for 20 min.

Bound antibody was detected using chemiluminescence (Pierce ECL Plus substrate, Thermo Fisher). The membrane was covered in substrate solution and placed between two clear plastic sheets before incremental imaging at 2 min intervals in a LAS-4000 camera. The intensity of the peptides in each membrane was measured using ImageJ and normalized to the controls (10xHis) and blank spots (100%-0%) using the the Protein Array Analyzer macro v1.1c^66^. For the CelluSpot arrays^67^, the intensity for each individual spot was also normalized by dividing each raw intensity with that of the equivalent spot in a membrane baited with His6-SUMO (produced in-house), without partner protein

### Biolayer interferometry (BLI)

Pre-hydrated streptavidin Octet Biosensors (FortéBio) were coated with 4 µM solutions of biotinylated HMGB1 proteins in 10 mM HEPES, pH 7.5, 150 mM NaCl (Base buffer, BB) plus, 0.5 mM TCEP (60 second baseline, 60 second binding). Nonspecific binding was minimized by incubation for 3 min in BB + 1% BSA + 0.05% Tween-20 (Kinetics Buffer, KB) prior to kinetics assays. Interaction with CXCL12 was measured using OctetRed 384 by incremental immersion of the sensors in solutions with increasing CXCL12 concentration (0-150 µM, in 1:2 dilutions) in KB (60 second baseline, 500 second association, 420 second dissociation, 180 second reduction in BB + 0.5 mM TCEP). Kinetics data were extracted using DataAnalysis v9.0 (FortéBio). Response at equilibrium (*R_Eq_)* was plotted against concentration in a Michaelis-Menten saturation plot to calculate *kD/B_max_* (saturation plot of concentration against *rEq*). Kinetic rates (association rate; *k_on_* and dissociation rate; *k_off_)* were derived from the direct measurements of each parameter from all the association and dissociation steps in the interferogram and fitted to a horizontal line (mean) across all measurements. Data from each replicate run were pooled in the same manner to calculate the overall mean. Fitting was modeled as a 1:1 binding mechanism on the assumption that CXCL12 did not have cooperative binding across HMG Boxes. Comparisons of parameters across curve fits were initially performed via Aikaike’s Information Criterion (corrected) (AICc) directly from the curve fit data. If a model with different kinetic constants was supported, these were compared by either Brown-Forsythe one-way ANOVA (*k_On_, k_Off_*) or 2-way ANOVA if the same value was calculated via multiple methods (*K_d_*), to assess whether both calculations were equivalent.

### Nuclear magnetic resonance (NMR) spectroscopy

^15^N-labelled recombinant HMGB1 in 10 mM HEPES, 150 mM NaCl, pH 7.5 (matching BLI buffer conditions) was supplemented with 5% D₂O and loaded into 5 mm Shigemi tubes (>330 µL final volume, sealed with parafilm). CXCL12, prepared in identical buffer, was titrated stepwise into the same NMR tube, and an HSQC spectrum was acquired after each addition. NMR data were collected using spectrometers operating at ^1^H frequencies of 500 and 750 MHz; both spectrometers are equipped with 5 mm TCI cryoprobes. The 500 and 750 MHz spectrometers are equipped with Avance II and Avance IIIHD consoles, respectively. Peaks in ^1^H-^15^N-HSQC spectra were assigned based on published NMR chemical shifts for HMGB1 1-184 (BMRB 15418) and using 3D ^15^N-edited NOESY and TOCSY spectra collected for each construct. CXCL12 binding was measured using the chemical shift and intensity changes observed for samples with different molar equivalences of CXCL12 as listed in the corresponding figures. Chemical shift perturbations (CSPs) were calculated using the standard combined equation:

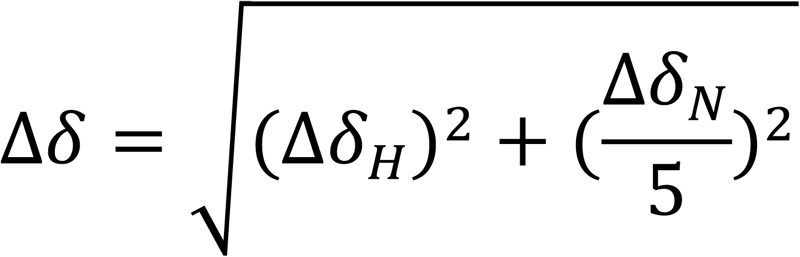

where Δ*δ_H_*and Δ*δ_N_*represent the observed changes in ^1^H and ^15^N chemical shifts, respectively. Spectra were analyzed using CCPNMR 3.0^68^. A sample of each HMGB1 construct was kept separate and measured after the titration experiment to account for chemical shift drifting during the experiment without CXCL12. For peak intensity changes, the median intensity change in each set was considered as a baseline. In the experiments with 3S-HMGB1 1-184, saturation to the same level as with the HMG Boxes alone could not be achieved due to solubility constraints. Box B constructs or 3S-HMGB1 1-184 were used to preclude the need for reducing agents to maintain the HMGB1 in the fully reduced state as these could also reduce the disulfide bonds of CXCL12.

### RAGE binding assays

Due to the lack of an established biological signaling assay for RAGE without interference from other NF-κβ signaling, we assessed binding to HMGB1 *in vitro.* 384-well protein-binding ELISA plates (Santa Cruz Biotechnology) were coated with 50 µL of 40 nM solutions in PBS (+ 0.5 mM TCEP for FR-HMGB1 constructs) of HMGB1 constructs for 24 h at 4°C, including full-length DS-HMGB1 controls and blank, with 4 replicates of each. Nonspecific binding was blocked by incubation with 10% BSA in PBS for 2 h at 20-25°C. RAGE-Fc chimera protein (BioTechne; 0-640 nM in 1:4 dilutions) was added in 10% BSA/PBS and allowed to bind for 2 h at 4°C. Bound Fc chimera was detected by incubation with anti-Human IgG HRP (Agilent Dako) diluted 1:10000 in 1% BSA/PBS for 2 h at 20-25°C. The plate was washed 3 times with 100 µL PBST between each of these 3 steps. 25 µL of TMB substrate (Thermo Fisher) was added to each well to detect bound antibody. The reaction was allowed to develop in the dark until the full-length DS-HMGB1 control developed a concentration-dependent color gradient, before stopping the reaction with 25 µL of 0.5 M H_2_SO_4_. OD450 was used as a readout (FluoStar OMEGA, BMG Labtech) and plotted as a saturation fit against 2x RAGE-Fc concentration as the chimera is a RAGE dimer.

RAGE binding kinetics to HMGB1 were measured by immobilizing 15 µg/mL RAGE-Fc in PBS + 0.1% BSA + 0.02% Tween-20 on the surface of anti-IgG biosensors (AHC) over 30 seconds and dipped in serial concentrations of each HMGB1 construct (60 second baseline, 200 second association/dissociation) and fitted in the same manner to derive kinetic parameters. All other parameters were identical to those for Biolayer interferometry with HMGB1.

### TLR-4 and TLR-2-mediated NF-κB signaling reporter assay

HEK-Dual cells (InvivoGen) expressing human TLR-2 and CD14 or murine TLR-4, MD-2 and CD14 were maintained in DMEM (Gibco), supplemented with 10% FBS (Gibco), 1% L-Glutamine (Gibco) and 1% penicillin/streptomycin (Gibco), in standard tissue culture conditions (37°C; 5% CO_2_). FR-HMGB1, DS-HMGB1 or dBB12L induced activation of TLR-4 and TLR-2 signaling was determined by plating 10^4^ TLR-4 and TLR-2 HEK-Dual cells in triplicate into 96 well plate and stimulating with 10 µg/mL HMGB1 and 100 ng/mL FSL-1 (Tocris Bioscience) for TLR-2 and 10 ng/mL LPS (Sigma-Aldrich) for TLR-4. NF-κβ activity was determined 24 h after stimulation by measuring the levels of secreted embryonic alkaline phosphatase (SEAP).

### Monocyte total TNF secretion assay

Human monocytes (Stem Cell Technologies) were maintained in DMEM (Gibco), supplemented with 10% FBS (Gibco) in standard tissue culture conditions (37°C; 5% CO_2_). Proinflammatory cytokine production by FR-HMGB1, DS-HMGB1 or dBB12L was determined by plating 10^5^ human monocytes in triplicate into 96-well plate and stimulating with 10 µg/mL HMGB1 and 50 ng/mL LPS (Sigma-Aldrich) or 10 ng/mL LTA (InvivoGen). TNF levels were determined 24 h after stimulation using Enzyme-Linked Immunosorbent Assay (ELISA) (Abcam).

### *In vivo* mouse muscle injury model

Procedures were performed as described previously^10,16^. Animals were anesthetized using aerosolized 2% isoflurane, given analgesia, transferred to a warming pad and the right lower hind limb was disinfected with povidone iodine and the tail with 70% ethanol if intravenous injection was performed. 50 μL of 1.2% BaCl_2_ (Sigma-Aldrich) was injected along the length of the tibialis anterior (TA) muscle to induce injury. Mice were injected with HMGB1 constructs (1.5 mg/kg, 56 nmol/kg, resuspended in PBS) or PBS vehicle control intravenously at the time of injury or later to determine efficacy of delayed administration.

Mice were euthanized and lower limbs removed at the times indicated and fixed in 4% paraformaldehyde (Santa Cruz Biotechnology) for 24 h. The TA muscles were dissected and fixed for a further 24 h before embedding in paraffin. Sections (5 µm) were stained with hematoxylin and eosin and imaged with an Olympus BX51 using a 10x ocular/ 40x objective lens. Myofibers are identified as polygonal, eosinophilic structures with distinct boundaries, containing either peripheral (mature) or centrally located (regenerating) nuclei. The cross-sectional area (CSA) of the fibers from at least 4 images per mice was manually measured using the FIJI distribution of ImageJ2 software (NIH).

### hMSC Osteogenesis Screen

Human bone marrow derived mesenchymal stem cells (hMSCs, Zenbio) were maintained in DMEM (Gibco) supplemented with 10% FBS (Gibco), 1% L-glutamine (GE Healthcare) and 1% penicillin/streptomycin (GE Healthcare) in standard tissue-culture conditions (37 °C; 5% CO_2_) and used between passages 3–5. The effect of dBB12L and FR-HMGB1 on osteogenesis were assessed by adding 10 µg/mL in 200 μL to 10^4^ hMSCs cultured in maintenance media and plated in triplicate in a 96-well plate. Anti-CXCL12 neutralizing antibody (10 µg/mL;R&D Systems), glycyrrhizin (GL, 100 µM; Sigma-Aldrich) or CXCR4 antagonist AMD3100 (1 µM; Tocris Bioscience) were added 15 min prior to HMGB1 and maintained in the medium with every medium change. After 16 h, this was changed to osteogenic medium alone comprising maintenance medium supplemented with 100 nM dexamethasone, 50 μg/mL ascorbic acid 2-phosphate, and 10 mM β-glycerophosphate (all Sigma-Aldrich). The media was replaced at day 3, and at day 7 it was removed and the cells were lysed to assess the alkaline phosphatase activity, a marker of osteogenic differentiation, using commercial assay kits (Abcam).

### C2C12 myogenic differentiation assay

C2C12 cells were cultured in DMEM (Gibco) supplemented with 10% FBS (Gibco), 1% L-glutamine (GE Healthcare), and 1% penicillin/streptomycin (GE Healthcare) under standard tissue culture conditions (37 °C, 5% CO₂). Cells were seeded at 2 × 10⁴ cells/cm² and allowed to adhere overnight before initiation of the assay. FR-HMGB1 or dBB12L (1 µM) was added 16 h prior to induction of differentiation. Anti-CXCL12 neutralizing antibody (10 µg/mL; R&D Systems), GL (100 µM; Sigma-Aldrich), or the CXCR4 antagonist AMD3100 (1 µM; Tocris Bioscience) was added 15 min before HMGB1 treatment and maintained in the culture medium throughout the experiment, including during each medium change. To induce differentiation, growth medium was replaced with differentiation medium consisting of DMEM (Gibco) supplemented with 5% normal horse serum (Gibco), 1% L-glutamine (GE Healthcare), and 1% penicillin/streptomycin (GE Healthcare). After 2 days of differentiation, total RNA was isolated using the RNeasy Kit (Qiagen). One microgram of RNA was reverse transcribed using SuperScript IV (Thermo Fisher Scientific). The resulting cDNA was diluted 10-fold prior to SYBR Green–based quantitative PCR analysis. The primer sequences for *Myhc* were: forward, ACCGAAGGCGGAACTACTGTAAC; reverse, ATCCAGGCTGCATAACGCTCT. *Gapdh* was used as the housekeeping gene (forward, AGGTCGGTGTGAACGGATTTG; reverse, TGTAGACCATGTAGTTGAGGTCA). Results are presented as fold change relative to the untreated control (no drug added).

### HMGB1 ELISA for Mouse Serum Samples

Female C57BL/6J mice (10–11 weeks old) were injected with dBB12L. At designated time points, mice were euthanized and blood samples collected via cardiac puncture. The blood was allowed to clot at room temperature for 30 minutes and then centrifuged at 2,000 × g for 10 minutes at 4 °C to separate the serum. The serum was aliquoted and stored at −80 °C until analysis. For dBB12L detection, Nunc MaxiSorp 96-well plates (Thermo Fisher Scientific) were coated overnight at 4 °C with 50 µL per well of anti–HMGB1 antibody (Abcam) at a concentration of 1 µg/mL in PBS. The next day, plates were washed three times with PBS containing 0.05% Tween-20 (PBST). Wells were blocked with 100 µL per well of SOMAmer blocking buffer (containing 150 mM NaCl, 40 mM HEPES, 5 mM MgCl₂, 1 mM EDTA, 1% BSA (Sigma-Aldrich), 0.5 mM dextran sulfate, and 1 µM Z-Block (Standard BioTools) for 1 h at room temperature, followed by three washes with PBST. A standard curve was generated using dBB12L (stock concentration 1.4 mg/mL), serially diluted 1:3 starting from 200 ng/mL to 0 ng/mL in SOMAmer blocking buffer supplemented with 0.5 µM Z-Block. Standards were incubated overnight at 4 °C, then washed three times with PBST. A biotinylated oligonucleotide probe generated against dBB12L (Standard BioTools) was heat-denatured at 95 °C for 10 minutes and allowed to cool to room temperature. It was then diluted to 5 nM in SOMAmer blocking buffer containing 0.5 µM Z-Block and incubated on the plate for 3 h at room temperature. Plates were again washed three times with PBST. Detection was performed using streptavidin-HRP (R&D Systems) diluted 1:400 in buffer containing 0.5 µM Z-Block and 0.5% BSA. The detection reagent was incubated for 1 h at room temperature, followed by three washes with PBST. The signal was developed using 50 µL per well of TMB substrate (Thermo Fisher Scientific), and the reaction was stopped with 50 µL of 1 M sulfuric acid. Absorbance was measured at 450 nm using a microplate reader.

## QUANTIFICATION AND STATISTICAL ANALYSIS

All statistical analyses were performed using GraphPad Prism (v. 8.41). For kinetics experiments (BLI/RAGE ELISA), all fits were performed via nonlinear least squares. For the RAGE ELISA, as each concentration of RAGE was independent from the rest of the wells, all data were considered as one kinetics fit. For BLI, each sensor was considered an independent fit for the purposes of calculation. Comparisons between parameters were performed using the AUC method. Mouse muscle injury model data were analyzed using nested ANOVA, where each sub-column comprised all the muscle CSA values for a given animal, and each group contained all the animals in order to separate biological variation from treatment effect. If the equal variance assumption could not be met in either case, data were analyzed using a Kruskal-Wallis test; for nested ANOVA equal numbers of data from each animal were randomly selected to avoid skewing. Other data were analyzed using one-way ANOVA if the heteroscedasticity plot and Q/Q plot supported the equal variance assumption^69^, and verified using Spearmańs test. Post-hoc comparisons were weighted by Holms-Sidak correction (ANOVA family tests) or Dunńs method (Kruskal-Wallis). Data are presented as mean ± SEM. Each number represents a biological sample, and each sample with 3 technical replicates. Significance is indicated as *P < 0.05, **P < 0.01, ***P < 0.001, ****P < 0.0001.

### Declare of Interest

AVG, CL, AIES, N B-B, WY and JN are inventors on a patent application filed by Oxford University Innovation. JN is a co-founder of Salamander Therapeutics Corp.

## Acknowledgments

The study was supported by the Wellcome Trust (AZR02170).

