## Supplemental Information for "Engineered HMGB1 construct with tandem Box B domains promotes tissue regeneration without potential for inflammation"

**This file includes:**

Figure S1 to S8

Figure legends S1 to S8

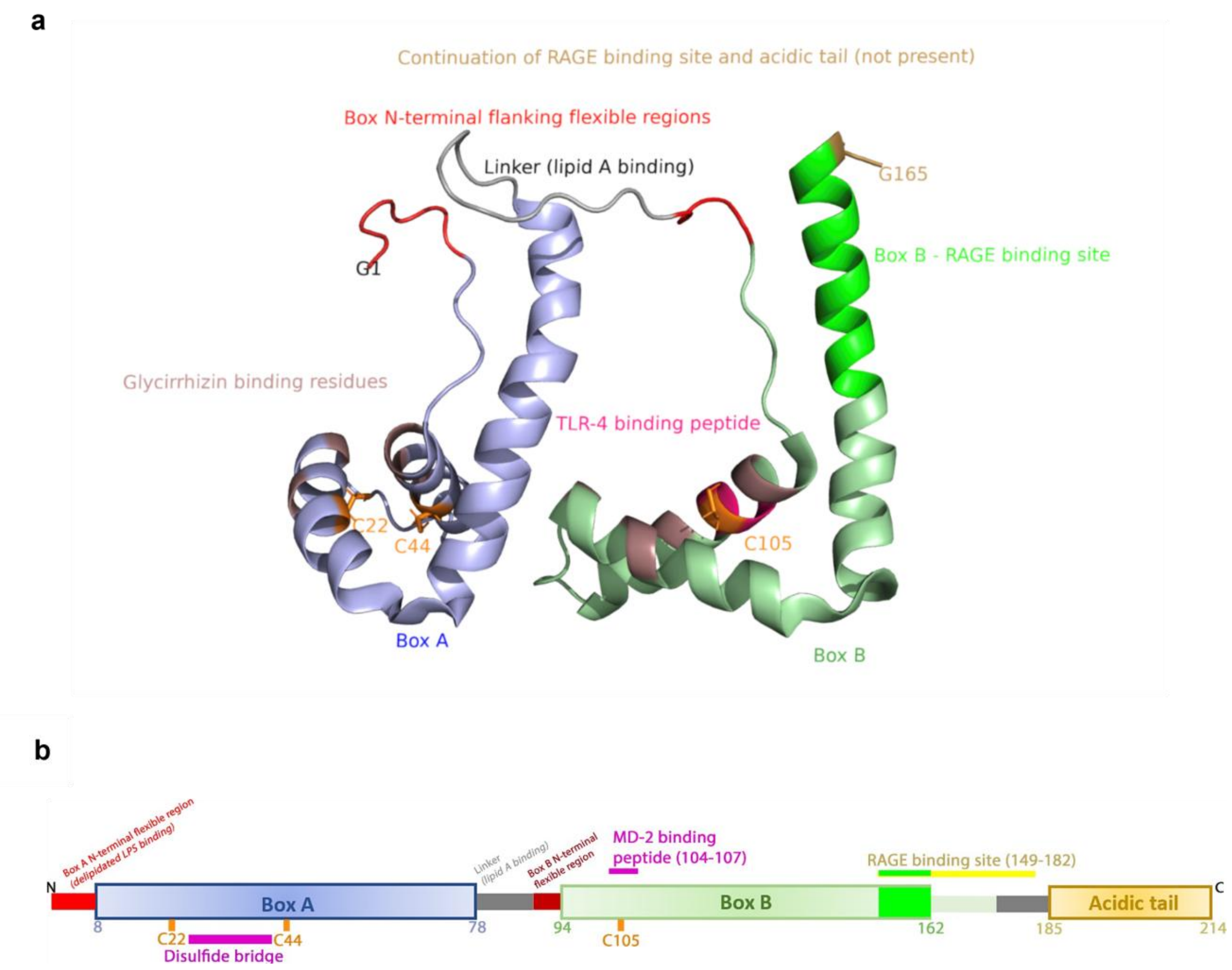


**Figure S1: Schematic of HMGB1 structures and known binding regions for RAGE and TLR-4. (a)** Structure of HMGB1 (PDB 2YRQ, conformer 1) with regions colored according to known interactions with LPS, TLR-4 and RAGE. The regions involved in TLR-2 binding have not previously been described. The acidic tail is not shown in the PDB structure. Pink: residues involved in glycyrrhizin binding. Red: flexible N- terminal regions adjacent to Box A or Box B. Orange: cysteine residues. White: linker region between HMG Boxes. Bright green and yellow: RAGE binding regions. **(b)** Schematic representation of regions within HMGB1 known to bind TLR-4/MD2 and RAGE.

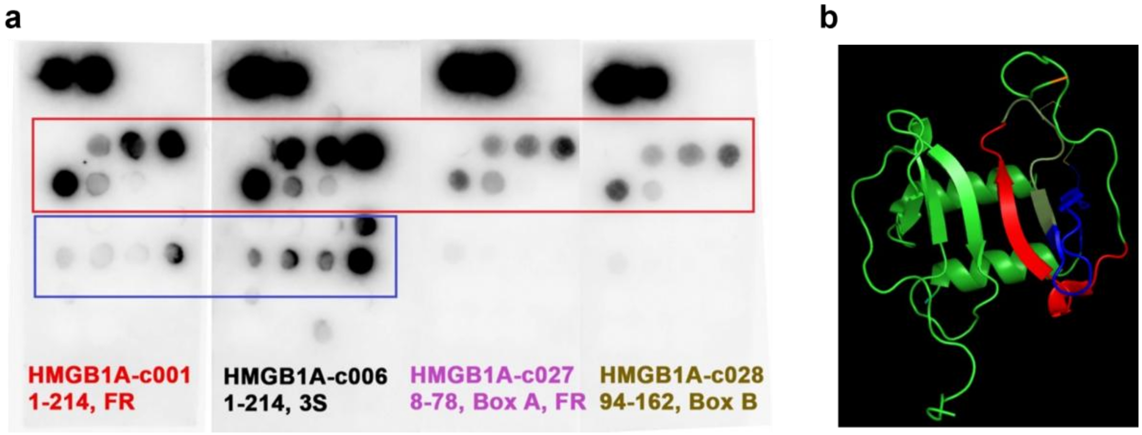


**Figure S2: Peptide array of CXCL12 peptides interacting with HMGB1. (a)** Peptide binding profiles show that full-length HMGB1 interacts with multiple regions of CXCL12, whereas individual Box A or Box B domains alone fail to bind certain peptides (highlighted in blue), indicating that the structural core of HMG Box A (8-78) and Box B (94-162) alone is not sufficient for full interaction with CXCL12 peptides. A common CXCL12-binding peptide (highlighted in red) exhibits reduced signal intensity when probed with Box domains alone compared to full-length HMGB1, confirming cooperative interaction. The array was probed with HMGB1 (FR or 3S)-His6 (1-214), Box A-His6 (8-78) or Box B-His6 (94-162). **(b)** Structural model of the CXCL12 dimer (PDB 2J7Z) with HMGB1-interacting regions highlighted: red, shared binding region detected with both full-length FR- or 3S-HMGB1 and individual Box domains; blue, non-shared regions bound only by full-length FR- or 3S-HMGB1, indicating a requirement for domain cooperation.

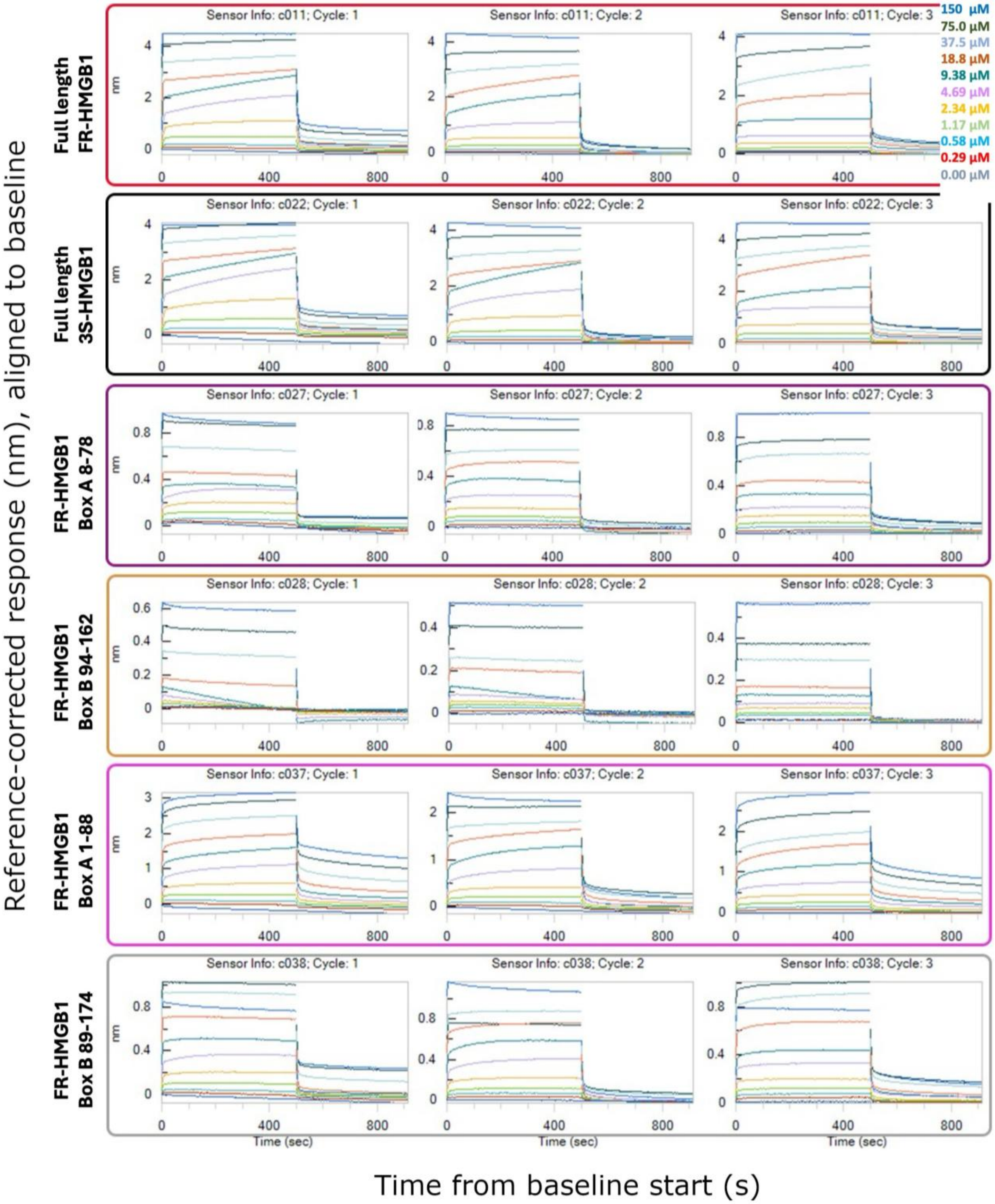


**Figure S3: BLI interferograms of CXCL12 binding to immobilized HMGB1 constructs.** Interferograms showing the reference-corrected response in nm (y-axis) over time in seconds (x-axis) of CXCL12 binding to biotinylated HMGB1 constructs colored according to CXCL12 concentration (key top right). Each set of three replicates (cycles) for a given sensor surrounded by a colored overlay according to construct.

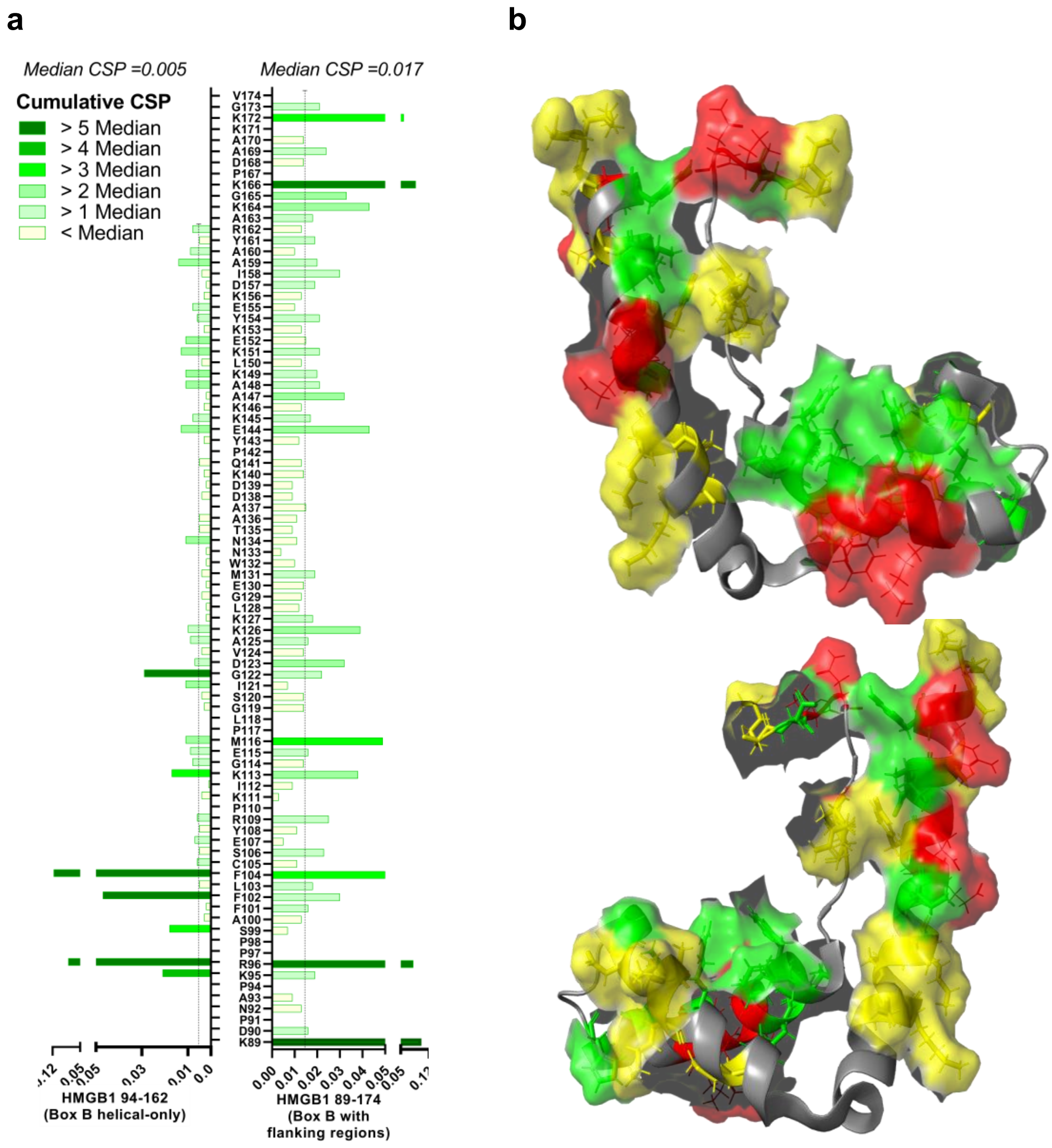

**Figure S4: NMR validation of CXCL12 binding interface within Box B. (a)** Cumulative CSP of biotinylated Box B constructs (FR 94-162 and FR-89-174) after titration with CXCL12 (0.42, 0.84 and 1.42 molar equivalents). The sequence of each HMGB1 construct has been overlaid with residue numbers; rows with no numbers represent residues which could not be assigned unambiguously in the HSQC spectra. Residues within CXCL12 binding peptides (underlined), not assessable by alanine scan, shown in gray. Dashed lines indicate median CSP. **(b)** Structure of FR-HMGB1 box B 89-166 (PDB: 2YRQ), with residues and their side chains colored according to the data (Two angles are shown). Red: residues identified in the peptide array as binding CXCL12. Yellow: Residues identified in NMR. Green: Residues identified in both NMR and peptide array.

a

b

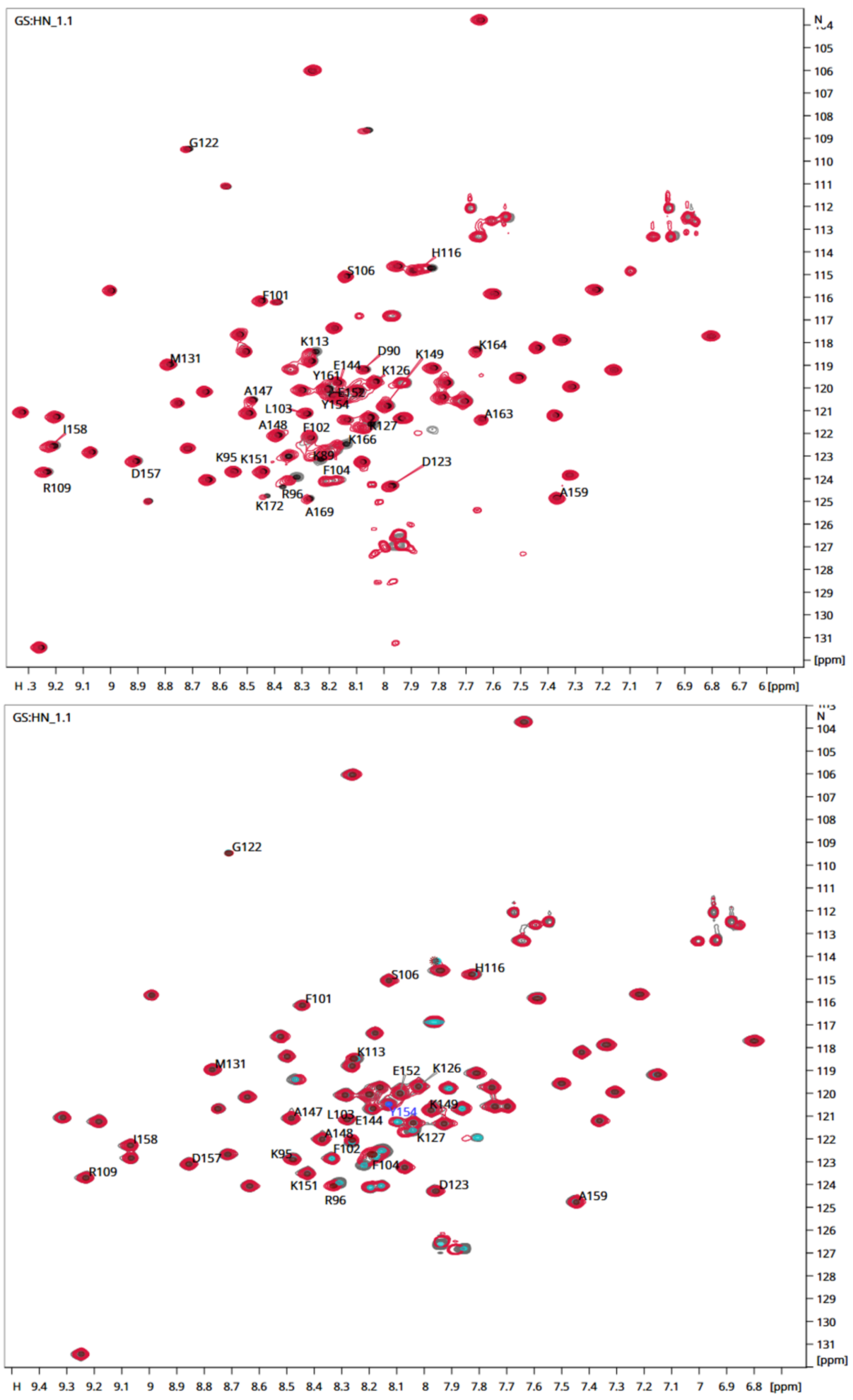


**Figure S5: Overlaid 500 MHz $^{15}N$ HSQC spectra of HMGB1 Box B constructs in 10 mM HEPES 150 mM NaCl pH 7.5 buffer. (a)** Overlay of FR 89-174: complete Box B (gray) and FR89-174 with CXCL12 in a 1:1.42 ratio (red). **(b)** Overlay of FR 94-162: helical-only biotinylated Box B (gray) and FR 94-162 with CXCL12 in a 1:1.42 ratio (red). Calculated CSPs are shown in Figure S4a. Selected residues exhibiting the largest chemical shift perturbations are indicated and labelled with residue numbers.

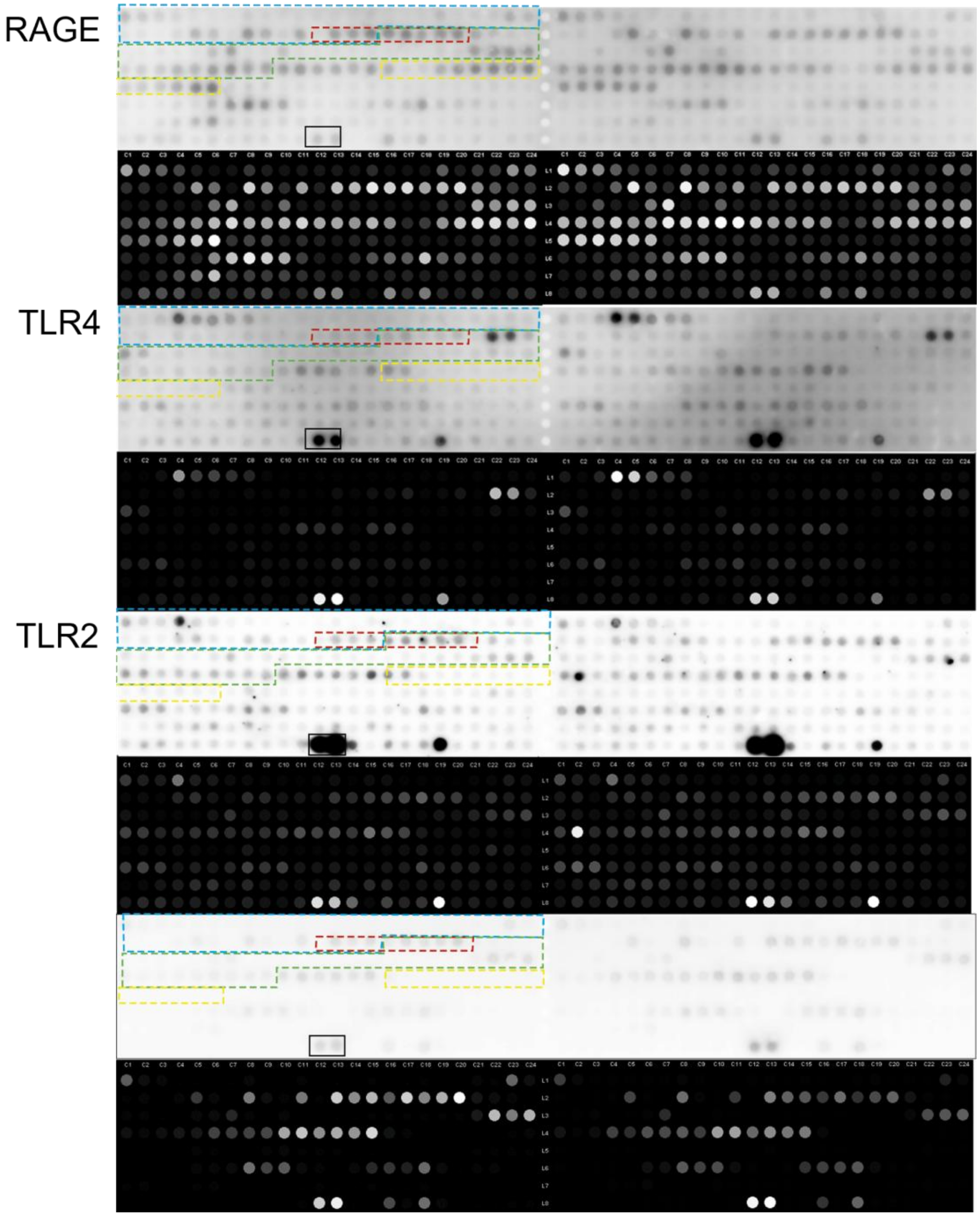


**Figure S6: DAMP receptor peptide arrays.** CelluSpot peptide arrays of HMGB1 peptides against RAGE, TLR-4 or TLR-2 and His6-SUMO control. For each set, top (grayscale) image shows the raw data from each membrane: bottom image (black with white dots) represents the intensities used for the calculation. Dotted lines represent the regions of HMGB1 whose sequences are contained in the spots. Blue, Box A; Red, linker; Green, Box B; Yellow, acidic tail.

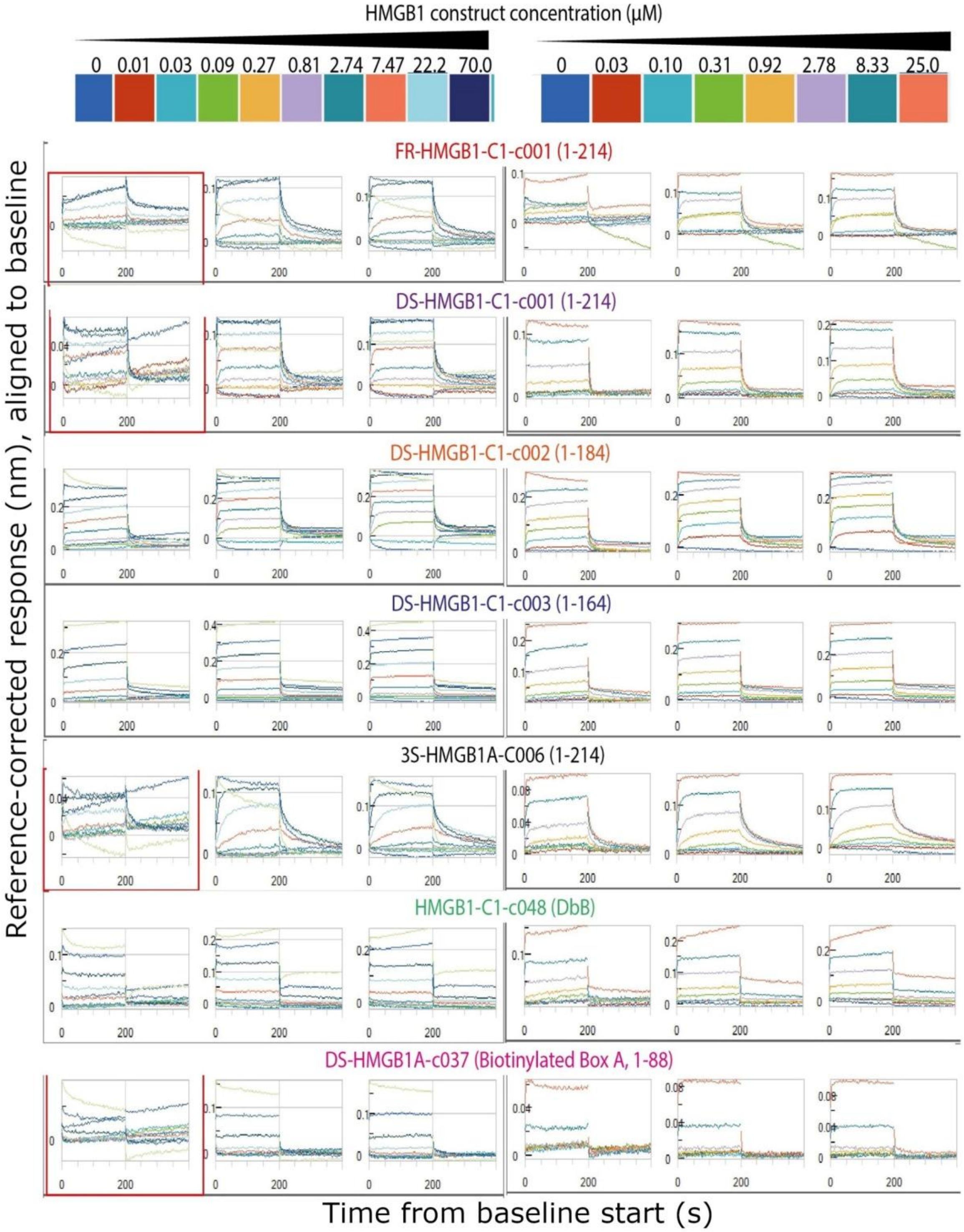


**Figure S7: Interferograms in BLI of binding of HMGB1 constructs to immobilized Fc-RAGE.** Left (first three columns of interferograms): 0 - 22.22 μM HMGB1 over 9 steps; right (next three columns of interferograms): 0 - 25 μM over 7 steps. Individual curves in each interferogram color-coded by concentration (top). Data for each graph was generated using a single sensor (replicate). Interferograms with data points excluded due to poor quality (e.g. drift) surrounded by a red rectangle. For all interferograms, the x-axis represents time in seconds, and the y-axis represents the reference-corrected response in nm.

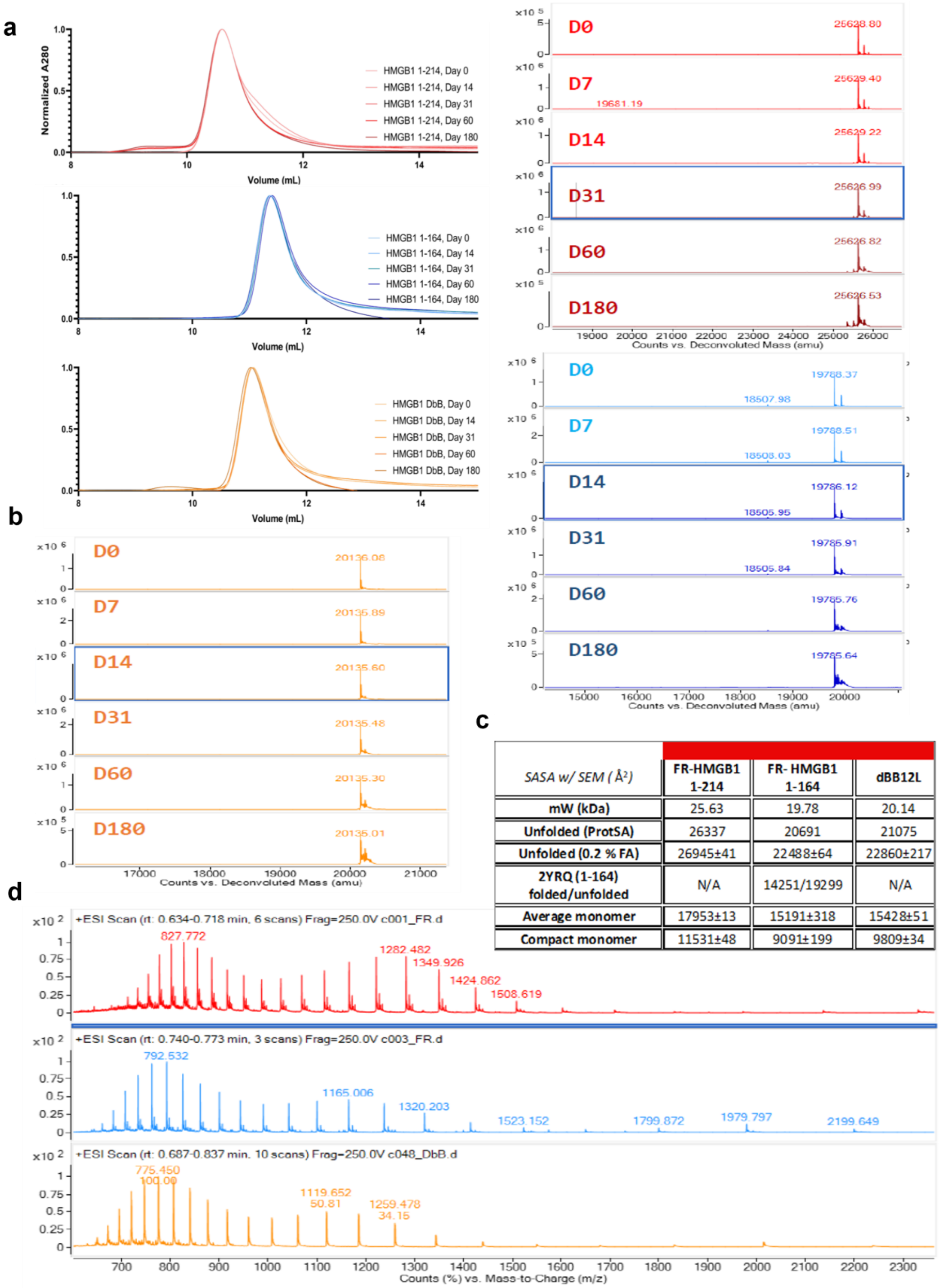


| SASA w/ SEM (Å²) | FR-HMGB1 1-214 | FR- HMGB1 1-164 | dBB12L |
|---|---|---|---|
| mW (kDa) | 25.63 | 19.78 | 20.14 |
| Unfolded (ProtSA) | 26337 | 20691 | 21075 |
| Unfolded (0.2 % FA) | 26945±41 | 22488±64 | 22860±217 |
| 2YRQ (1-164) folded/unfolded | N/A | 14251/19299 | N/A |
| Average monomer | 17953±13 | 15191±318 | 15428±51 |
| Compact monomer | 11531±48 | 9091±199 | 9809±34 |

**Figure S8 SEC of HMGB1 samples up to 180 days and parallel denaturing ESI-MS validation.** **(a)** Analytical SEC (Superdex S75 10/300 GL) of HMGB1 constructs; full length FR-HMGB1 (red), 1-164 FR-HMGB1 (blue) or dBB12L (orange) at days 0, 7, 14, 31, 60 and 180 after purification and endotoxin removal. Protein stored at room temperature in PBS with 2 mM TCEP. **(b)** Deconvoluted ESI-MS spectra showing no change in molecular weight of full-length FR-HMGB1, 1-164 FR-HMGB1 and dBB12L (lines colour-coded as panel a) as a result of degradation up to day 60; most of the protein is still intact at that time, with oxidation observed from day 31. **(c)** A summary of Retention volume in SEC showing average, high M/Z distribution alone, unfolded peptide in denaturing conditions, theoretical unfolded SASA (ProtSA), and the for the published structure of FR HMGB1 1-164 (PDB 2YRQ). **(d)** Sample denaturing M/Z spectrum for the unfolded SASA calculations.
